# Uncovering network mechanism underlying thalamic deep brain stimulation

**DOI:** 10.1101/2023.12.09.570924

**Authors:** Yupeng Tian, Edward Bello, David Crompton, Zoe Paraskevopoulos, Suneil K. Kalia, Mojgan Hodaie, Andres M. Lozano, William D. Hutchison, Matthew D. Johnson, Milos R. Popovic, Luka Milosevic, Milad Lankarany

## Abstract

Thalamic ventral intermediate nucleus (Vim) is the primary surgical target of deep brain stimulation (DBS) for reducing symptoms of essential tremor. High-frequency Vim-DBS (≥100Hz) has been clinically effective, generating two experimentally-observed features in Vim spiking activity: 1) a large transient excitatory response (lasting <1s), followed by 2) a suppressed steady-state consisting of oscillations. Yet, mechanisms underlying these observations have not been fully understood by previous studies. In this work, we developed a network rate model and a novel parameter optimization method that accurately fit in-vivo single-unit recordings of Vim in human patients with essential tremor receiving a wide range of DBS frequencies (5∼200Hz). Our model incorporates both the DBS-induced synaptic plasticity of Vim neurons, and the recurrent connections among excitatory and inhibitory neurons in Vim-network. We hypothesized that besides inducing synaptic depression, the therapeutic mechanism of high-frequency Vim-DBS could be to engage more inhibitory neurons in stabilizing the underlying circuits.

**Graphical abstract:** 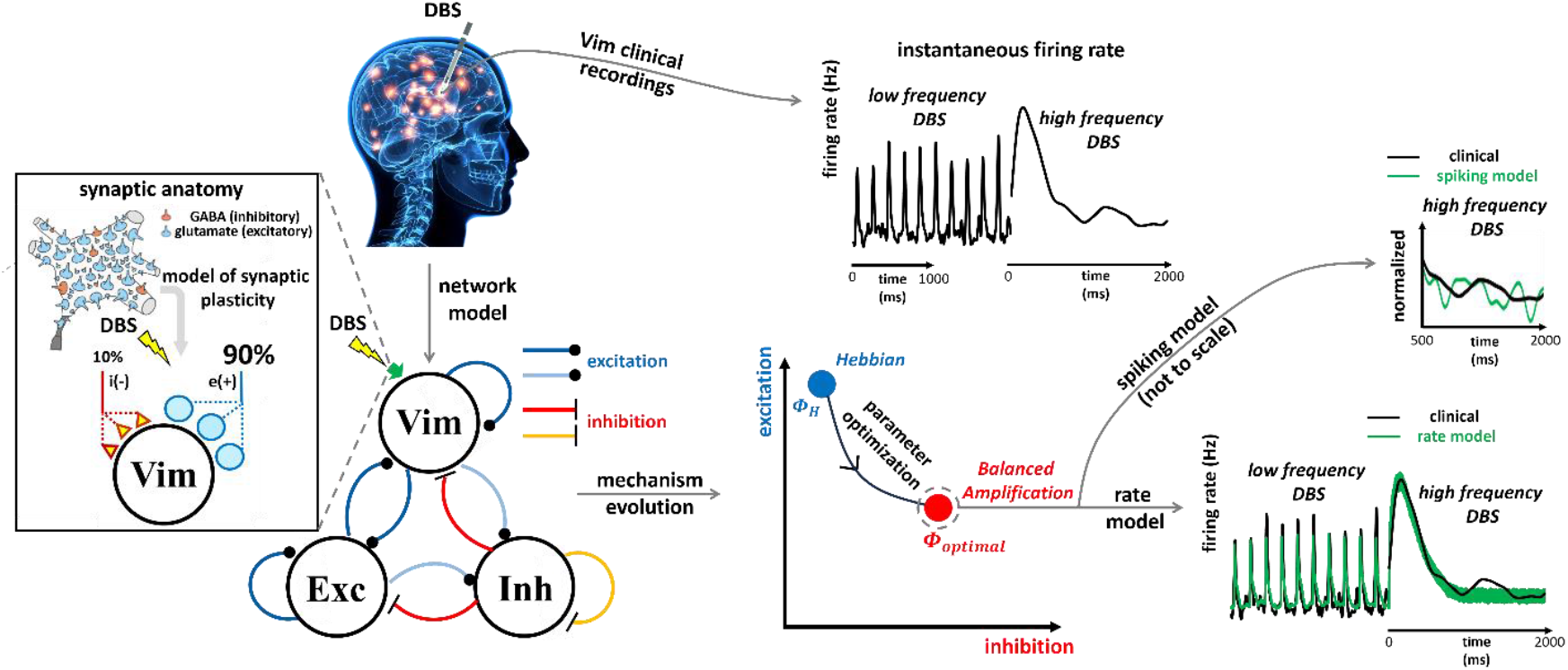

## Introduction

Deep brain stimulation (DBS) delivers electrical pulses to adjacent neuronal circuits and is known to modulate neuronal activity ^1,2,3^. DBS has become a standard therapy for many movement disorders, including Parkinson’s disease ^2^, essential tremor ^3^, and dystonia ^4^. DBS is now being investigated as a treatment for psychiatric or cognitive disorders, including depression ^5^, obsessive-compulsive disorder ^6^ and Alzheimer’s disease ^7^. DBS of the thalamic ventral intermediate nucleus (Vim) – i.e., Vim-DBS – is the primary surgical option of DBS for treating essential tremor ^8^,^9^. Essential tremor is the most common adult tremor disorder affecting up to 1% of adults over 40 years of age, and features attention tremor and uncontrollable shaking of the affected body parts ^10,11^.

Despite the recognized clinical benefits of DBS, its therapeutic mechanisms on the disease-affected neuronal circuits are not fully understood ^1,12^. High-frequency DBS (≥100 Hz) can be clinically effective for relieving symptoms of Parkinson’s disease ^2^, essential tremor ^9^ and depression ^5^. Single-unit recordings of Vim neurons receiving high-frequency DBS demonstrated two critical features of firing rate dynamics: 1) an initial large excitatory transient response, lasting for <1 s, followed by 2) a suppressed steady-state consisting of oscillations ^12,13^. Suppression of local activity was suggested as a primary mechanism of high-frequency DBS ^14,15^, which could induce effects such as synaptic plasticity ^12^, axonal failure ^16^ or GABAergic activation ^17^. These DBS effects could depend on a combination of factors such as the stimulation sites ^12^ and the time-course of synaptic depression ^18^. Besides these existing hypotheses, computational models are needed to quantify and further explore the underlying physiological mechanisms ^19,13^. The dynamics of DBS-induced membrane and local field potentials (LFP) are commonly modeled ^20,21,22^; yet, these models lack the efficacy of tracking instantaneous firing rate observed from single-unit recordings. A recently developed firing rate model of DBS-induced synaptic plasticity represented experimental single-unit recordings, but deviated from observations during high-frequency Vim-DBS ^13^. Thus, there is a need for a firing rate model of more detailed network mechanisms, besides synaptic plasticity. To the best of our knowledge, there has been no neuronal network model that can accurately track the instantaneous firing rate from in-vivo human single-unit recordings during DBS.

In a neuronal network, besides synaptic plasticity, the recurrent interplays among neurons are critical in forming the network dynamics ^23,24^. Feedback inhibition characterizes the recurrent connections between excitatory and inhibitory neurons, and is critical in stabilizing the neuronal networks, e.g., hippocampus ^25^, basal ganglia ^26^, sensory cortex ^27^ and thalamus ^28^. Lack of inhibitory effect in thalamic circuits could lead to excessive firings of thalamocortical relay neurons, whose over-activity could further induce essential tremor through corticomuscular projections ^29,30^. Murphy and Miller (2009) ^31^ proposed a Balanced Amplification mechanism demonstrating strong inhibition in response to strong external drive of excitatory neurons. Such strong external inputs could be specified as intensive visual stimuli ^31^, strong injection current ^32^, high-frequency DBS ^33^, etc. In addition to balancing the effect of strong inputs, during weak inputs (e.g., low-frequency DBS), the inhibitory nuclei are less activated, and this leads to a supralinear increase of neuronal firing rate ^34^. Such inhibition-stabilization mechanisms are ubiquitous in cortical networks because of the prevalence of cortical inhibitory interneurons ^24,35^, and should be also universal in general brain networks, e.g. cortical-subcortical networks ^22,26,20^, when the role of inhibitory nuclei is significant.

In this work, we developed a firing rate model of the neuronal network of Vim impacted by Vim-DBS. In our model, the Vim-network consists of recurrent connections among three neuronal groups: DBS-targeted Vim neurons, external excitatory nuclei and inhibitory nuclei. For the Vim neurons, the DBS-induced short-term synaptic plasticity is characterized by the Tsodyks & Markram model ^36^. The external nuclei are mainly from thalamus, cerebellum and motor cortex ^37,38,39^. The incorporation of an external excitatory feedback component to the model may allow the model to capture the indirect effects of thalamic-DBS as they recurrently propagate through the motor control network, away from and back towards the Vim ^37,38^. An inhibitory feedback component may capture the contributions of the thalamic reticular nucleus (TRN), as well as intra-nucleus inhibitory interneurons ^39,12^. We then developed a novel parameter optimization method that accurately fits the model to clinical data recorded from human patients receiving DBS with varying stimulation frequencies (range 5 to 200 Hz). During the optimization process, we observed that the network mechanism evolves from Hebbian (dominant recurrent excitation) ^24,23^ to Balanced Amplification (equally strong excitation and feedback inhibition), which accurately reproduces the initial large transient response observed during high-frequency DBS. We further developed a spiking model of the membrane potential dynamics ^22,40^, and found that the Balanced Amplification spiking network could characterize the oscillations observed in the steady-state response during high-frequency DBS. Then, we further observed Vim membrane potential data and found that the inhibitory effect was observed as evoked hyperpolarized potential in 5 out of 19 Vim neurons receiving high-frequency DBS, yet less often observed during low-frequency DBS ^33^. From our results, we hypothesized that high-frequency DBS could engage more firings from inhibitory neurons, which stabilize the underlying network and lead to better therapeutic outcomes.

Our models and optimization method can be potentially extended to identify and study various brain neuronal circuits. Our rate model can be implemented to optimize the DBS frequency in a closed-loop control system ^41^ potentially used in clinics.

## Results

### Framework of rate model, optimization and mechanism analysis

We developed a firing rate model of the Vim-network in patients with essential tremor (**Fig. 1**). The model was fitted to the firing rate dynamics from experimental single-unit recordings of human Vim neurons receiving DBS with different frequencies – 5, 10, 20, 30, 50, 100, and 200 Hz ^12^. The model incorporates the short-term synaptic plasticity (STP) ^36^ of DBS-impacted Vim neurons, and their recurrent connections with the external excitatory and inhibitory nuclei (**Fig. 1**). To detect the optimal model parameters, we developed a novel optimization method that found the consistent model parameters that accurately replicate the experimental data across different DBS frequencies (**Fig. 1**). The optimization includes the sequential fittings to Focused Feature (more clinically effective high-frequency Vim-DBS data ^8^) and Stabilized Feature (balancing fitting accuracy across data from various DBS frequencies) (**Fig. 1**). During the optimization process, we observed that the modeled Vim-network evolves from an excitation-dominant mechanism (Hebbian) ^42^ to an inhibition-stabilized mechanism (Balanced Amplification) ^24^ (**Fig. 1**). The optimal modeled Vim-network is characterized by a Balanced Amplification mechanism with strong recurrent excitation stabilized by equally strong feedback inhibition (**Fig. 1**).

**Fig. 1.**
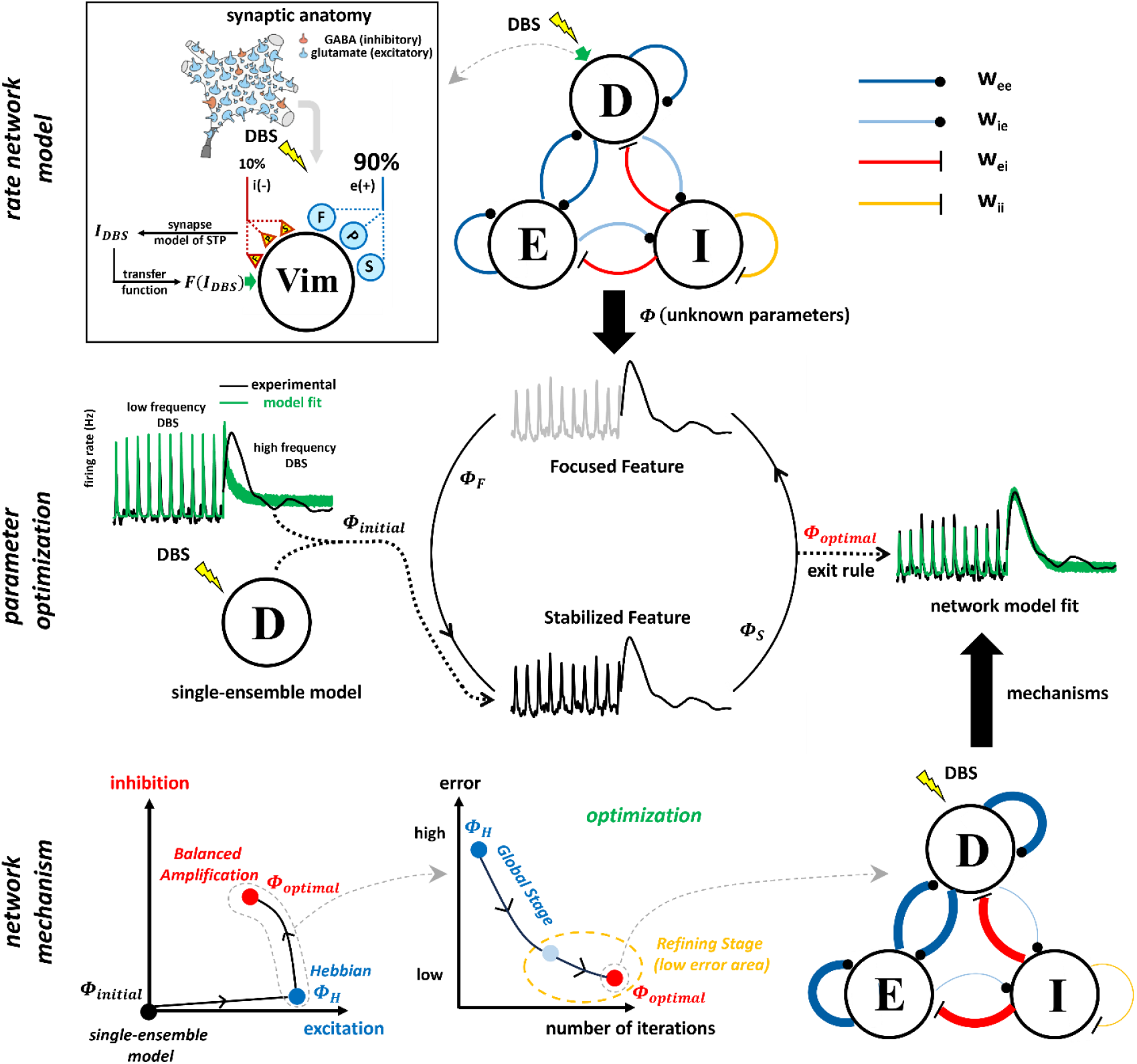
Schematic illustration of the rate model, optimization and mechanism analysis framework. (**rate network model**) The firing rate network model consists of 3 recurrent neural groups: “D” represents the ventral intermediate nucleus (Vim) neurons directly receiving DBS, “E” represents the external excitatory nuclei, and “I” represents the external inhibitory nuclei. w_ei_ represents the connectivity strength from inhibitory neurons to excitatory neurons; similar meanings for w_ee_, w_ie_ and w_ii_ . The connection with a dot (respectively, a bar) represents excitation (respectively, inhibition). DBS is delivered to the neural group D (Vim neurons), and we model the corresponding synaptic anatomic structure. Excitatory (respectively, inhibitory) synapses consist of 3 types: “F” (facilitation), “P” (pseudo-linear)”, and “S” (depression) (^12^, **Methods**). We formulate the DBS-induced post-synaptic current (I_DBS_) with the Tsodyks & Markram model of short-term synaptic plasticity (STP) (^36^, **Methods**). I_DBS_ is then transformed to the corresponding firing rate dynamics F(I_DBS_), which is the external DBS input into the rate network model. (**parameter optimization**) The model is fitted to human clinical Vim-DBS data. The initial model parameters (Φ_ini_) are from our previous work of a single-ensemble Vim model ^13^. We define two features of the clinical Vim-DBS data: Focused Feature stresses data from high frequency DBS, which is more clinically effective ^8^; Stabilized Feature balances the fitting accuracy between low and high frequency DBS data. During the optimization, the resulting parameters of fitting one feature are the initial parameters for fitting the other feature. The optimization loops are sequentially executed until an exit rule representing small fitting error is satisfied, and we obtain the optimal model parameters (Φ_optimal_). (**network mechanism**) During our optimization, we observe that the modeled network mechanism evolves from Hebbian (dominant excitation) to Balanced Amplification (inhibition comparable to excitation). The model fitting error fast decreases in optimization Global Stage (wide exploration of parameter space), and converges in Refining Stage (exploitation of the parameter rage with low fitting errors). The optimal network is characterized by a Balanced Amplification mechanism with equally strong excitation and inhibition.

### Firing rate network model

The rate network model of the Vim-DBS is illustrated in **Fig. 1**. The model consists of three differential equations,

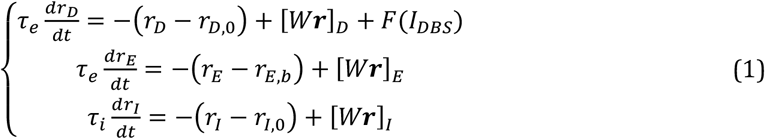

*where*,

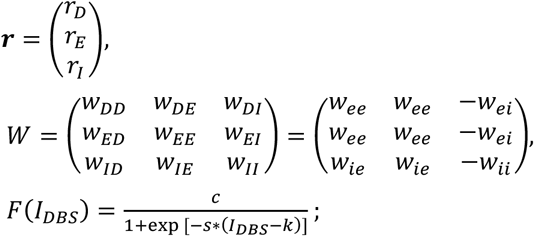

*The parameters* Φ = {*w*_*ee*_, *w*_*ie*,_ *w*_*ei*_, *w*_*ii*_, τ_*i*_, τ_*e*_, *r*_*E*,*b*_, *c, s, k*} *are undetermined*

In Equation (1), neural group “*D*” represents the Vim neurons directly receiving DBS, neural group “*E*” represents the external excitatory nuclei, and neural group “*I*” represents the external inhibitory nuclei (**Fig. 1**). The group “*E*” neurons are mainly the cerebellum dentate nucleus and the pyramidal cells of primary motor cortex (M1) deep layers (Layer 5 and 6) ^37,12,38^. The group “*I*” neurons mainly consist of the thalamic reticular nucleus (TRN) and interneurons ^39,12^. *r*_*m*_ (*m* ∈{*D, E, I*}) represents the firing rate of the corresponding neural group. The baseline firing rate (with DBS–OFF) of Vim neurons (*r*_*D*,0_) and the average firing rate of external inhibitory nuclei (*r*_*I*,0_) were obtained from single-unit recordings reported in other studies ^12,43^. We chose *r*_*D*,0_ = 25 Hz to be consistent with the human Vim experimental recordings in our previous work Milosevic et al. (2021) ^12^, and *r*_*I*,0_ = 5 Hz to be consistent with the experimental data recorded in both TRN (human) ^12^ and thalamic interneurons (mice) ^43^. Since the external excitatory nuclei (group “ *E* “) originate from multiple sources with highly variable firing rates ^44,45^, the corresponding baseline firing rate (*r*_*E*,*b*_) is left as an unknown variable (Equation (1)). In the experimental recordings from mice, the regular firing rate of cerebellum dentate nucleus ranged from 10 to 80 Hz ^44,46^. For M1 Layer 5 and 6 neurons in mice, the firing rate ranged from 10 to 60 Hz ^45,47^. Thus, we constrained *r*_*E*,*b*_ in the range of 10 to 70 Hz, and initialize it at *r*_*E*,*b*,0_ = 40 Hz.

In the rate network model, we use τ_*e*_ and τ_*i*_ to denote the excitatory and inhibitory time constants, respectively (Equation (1)). Since Vim neurons are excitatory ^12^, the time constant of the neural group “*D*” was τ_*e*_. In a firing rate model of a population of neurons, the time constant (τ) represents the changing speed of the firing rate in response to the post-synaptic current ^48^. Generally, time constants in firing rate models were considered in the range of 0 to 30 ms ^48,49^. The rate model time constant is consistent with the membrane time constant ^48^, and generally, the time constant of the inhibitory neurons is larger than that of the excitatory neurons ^12,50^.

The matrix W (Equation (1)) indicates strength of connectivity between different groups of neurons ^24,51^. In matrix W, *w*_*pq*_ (for *p*,*q* belong to group “*D*”, “*E*”, “*I* “) represents the connectivity strength from the neural group “*q*” to group “*p*”, and the +/-sign denotes the excitatory/inhibitory effect. The total network input into each neural group is computed as the matrix multiplication *W****r*** = ([*W****r***]_*D*_, [*W****r***]_*E*_, [*W****r***]_*I*_)^*T*^, where [*W****r***]_*D*_, [*W****r***]_*E*_ and [*W****r***]_*I*_ represent the inputs into group “*D*”, “*E*” and “*I*”, respectively (Equation (1)).

For the Vim neurons directly receiving DBS (group “*D*”), we modeled the DBS-induced post-synaptic current (*I*_*DBS*_) with the Tsodyks & Markram model ^36^ of short-term synaptic plasticity (STP) (**Methods, Supplementary Fig. S2**) in agreement with Milosevic et al. (2021)^12^; *I*_*DBS*_ is then transformed (with a sigmoid function) to the corresponding firing rate dynamics *F*(*I*_*DBS*_) (^13^, **Fig. 1, Supplementary Figs. S3 and S5**, Equation (1)). In the sigmoid transfer function *F*(*I*_*DBS*_), *c, s* and *k* are the scale, shape and shift parameters, respectively (Equation (1)).

The undetermined parameter set in the rate network model is *Φ* = {*w*_*ee*,_*w*_*ie*,_*w*_*ei*_, *w*_*ii*_, τ_*i*_, τ_*e*_, *r*_*E*,*b*_, *c, s, k*} (see Equation (1)). All rate model simulations were conducted with the sampling resolution of 0.1 ms.

### Proposed rate network model captures neural dynamics of human experimental data

The rate model was developed based on the experimental single-unit recordings of human Vim neurons in patients undergoing DBS surgery for essential tremor. While recording activity of an individual Vim neuron, DBS was applied with one of the stimulation frequencies {5, 10, 20, 30, 50, 100, and 200 Hz}, with specific stimulation length {10, 5, 3, 2, 1, 5, and 2 s}, respectively (see **Methods** for detailed data protocols which were already reported in our previous work ^12^). We obtained 5∼8 recordings during each frequency of DBS. During high-frequency DBS, we observed initial transient responses with intensive spikes; the transient response length (mean ± standard deviation) for 100-Hz and 200-Hz DBS is 690.76 ± 217.38 ms and 254.88 ± 59.88 ms, respectively (**Supplementary Table S10**). The transient response length during 100-Hz DBS is significantly longer than that during 200-Hz DBS (F (1,11) = 18.63, p = 0.0012, **Supplementary Table S10**), and this implies that the Vim spiking activity is suppressed to a higher extent during 200-Hz DBS. For data from each DBS frequency, we computed the instantaneous firing rate with a time histogram method based on these multiple recordings (**Methods**). The instantaneous firing rate was computed by convolving the experimentally recorded spike trains with an optimized Gaussian kernel that best characterized the spikes using a Poisson process (^52,53^, **Methods, Supplementary Fig. S1**). We concatenated the instantaneous firing rate of each DBS frequency, and optimized the consistent model parameters across different DBS frequencies (5 to 200 Hz) (^13^, **Supplementary Notes 1**). In **Fig. 2**, we showed the instantaneous firing rate calculated from experimental data, the results of our model fit, and the optimal model parameters.

**Fig. 2.**
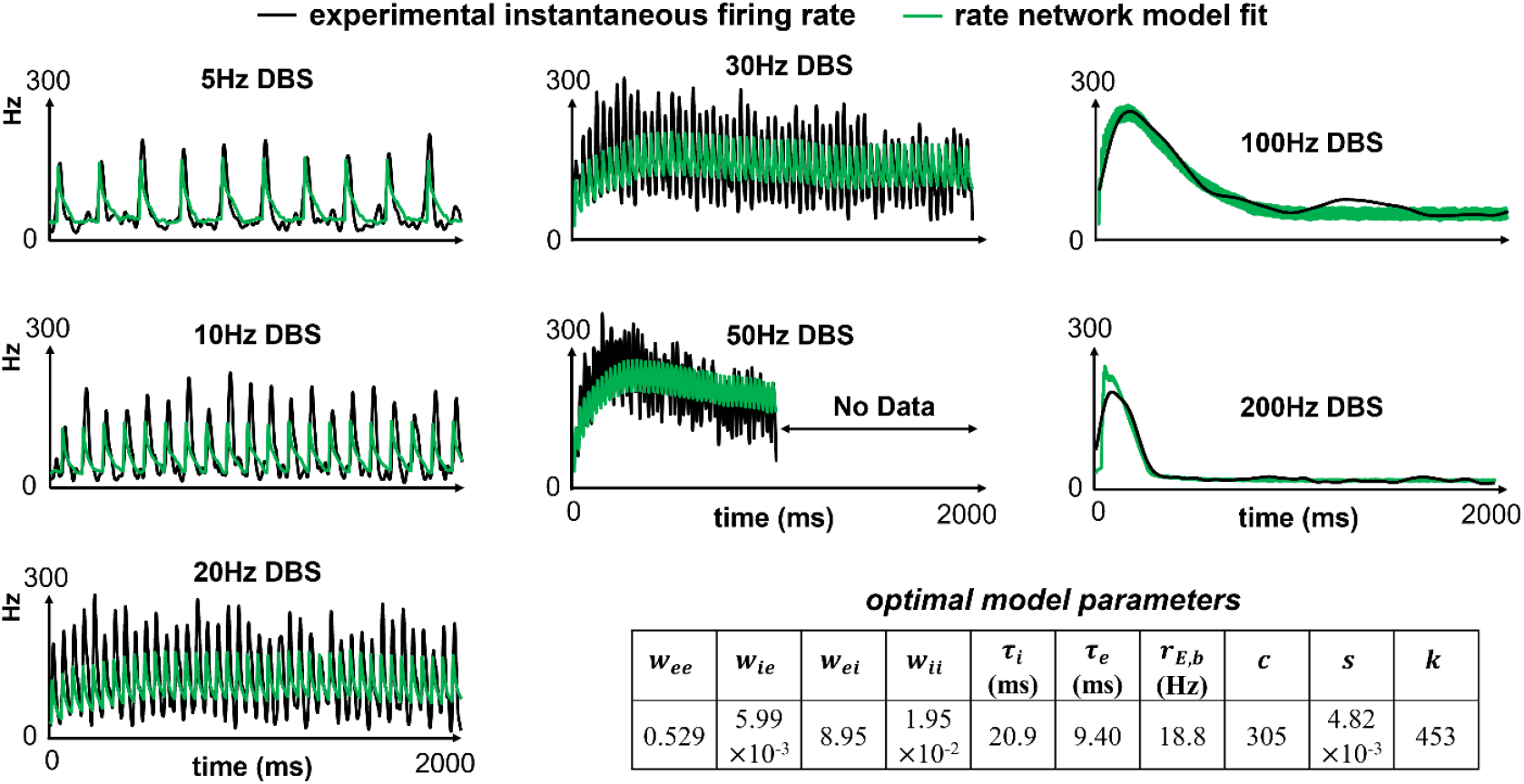
Fitting rate network model to Vim-DBS in humans. The firing rate network model is fitted to the experimental data recorded in the neurons in the ventral intermediate nucleus (Vim) of the human patients with essential tremor, during DBS of various stimulation frequencies (5 to 200 Hz). For each frequency of DBS, we obtained 5 to 8 spike trains from experimental single-unit recordings in different patients, and compute the instantaneous firing rate using a time histogram method. In computing the experimental instantaneous firing rate, we implement an optimized Gaussian kernel that best characterized the Poisson process underlying the spiking data (^53^, **Methods**). From the data of each DBS frequency, we present the model fit of the initial 2 seconds; an exception is 50-Hz DBS data, where the length of recording was ∼1 second. We compare the model fit (green line) with the experimental instantaneous firing rate (black line). The optimal model parameters (Equation (1)) are obtained with our route optimization method (see text). w_pq_ (p, q ∈{i, e}) is the connectivity strength (see **Fig. 1** legend for specific descriptions). τ_e_ and τ_i_ are the excitatory and inhibitory time constants, respectively. r_E,b_ is the baseline firing rate of the external excitatory nuclei (group E, **Fig. 1**). c, s and k are the phenomenological parameters in the sigmoid transfer function F (Equation (1)).

As shown in **Fig. 2**, the rate network model accurately reproduced the recorded firing rates across different DBS frequencies (5 to 200 Hz). The model could capture both transient and steady-state firing rate responses to each frequency of DBS. In particular, for 100-Hz and 200-Hz DBS data, the model is almost identical to the experimental data; this improves the result from our previous model that incorporated a single population of Vim neurons, and ignored the recurrent connections with other nuclei (^13^, **Supplementary Fig. S6**). For Vim-DBS, high-frequency DBS (100 to 200 Hz) is more clinically effective than low-frequency DBS (<100 Hz) ^8,9^. Additionally, we optimized the model parameters based on the concatenated signal across different DBS frequencies (5 to 200 Hz). When fitting a rate model to such DBS data with concatenated frequencies, the fit accuracy is consistent between observed and unobserved DBS frequencies (e.g., 130 Hz, 160 Hz; see Table 1 in Tian et al. (2023) ^13^).

We quantitatively validated the model goodness of fit by computing the normalized mean squared error (NMSE) between the experimental instantaneous firing rate (reference) and the model generated firing rate. Since high-frequency (100 to 200 Hz) Vim-DBS is more clinically effective ^8,9^ we emphasized the high-frequency DBS data, and defined the total fitting error (ER) for model validation:

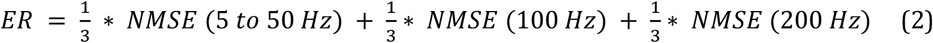

The NMSE (5 to 50 Hz) represents the NMSE of the model fit to the concatenated data from DBS frequencies 5 to 50 Hz. NMSE (100 Hz) and NMSE (200 Hz) were computed with data from the 100-Hz and 200-Hz DBS, respectively. For the rate network model fit shown in **Fig. 2**, ER = 7.6%, with NMSE (5 to 50 Hz) = 13.9%, NMSE (100 Hz) = 3.2% and NMSE (200 Hz) = 5.7%. See **Supplementary Table S7** for the NMSE of the model fit to data from each DBS frequency.

### Effective DBS-induced inputs characterize the balance between excitation and inhibition

The dynamics of a neural network is shaped by the effective transmission of the firing rate among interacting nuclei ^51,54^. Such transmission is determined by two factors: (i) the connectivity strength modeled by *W* in Equation (1); and (ii) the firing rate of the pre-synaptic nuclei. Thus, in our model, we formulate the effective input ^51^ into a neural group as the product of connectivity strength and pre-synaptic firing rate. The “effective input matrix” of the Vim-network receiving DBS is defined in Equation (3),

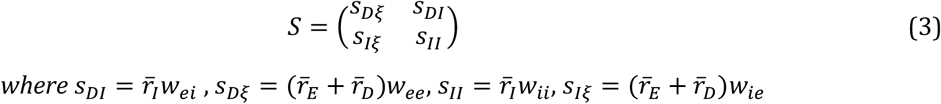

The subscript “*D*” represents the Vim neural group directly receiving DBS; “ *E* “ and “*I* “ represent the external excitatory and inhibitory nuclei, respectively (Equation (1), **Fig. 1**). The subscript “ξ” represents total input projected by excitatory neurons consisting of “*D*” and “*E*” groups of neurons. *s*_*mn*_ (*m* ∈{*D, I*}, *n* ∈{ξ, *I*}) indicates the effective input from group “*n*” to group “*m*”, and it is equal to the product of the corresponding connectivity strength and pre-synaptic firing rate. *w*_*pq*_ (*p, q* ∈{*e, i*}) is the modeled connectivity strength (Equation (1), **Fig. 1**). The pre-synaptic firing rate *r*_*n*_ (*n* ∈{*D, E, I*}) of the corresponding neural groups was calculated by averaging model generated firing rates across different DBS frequencies (5 to 200 Hz). See **Fig. 2** and **Supplementary Figs. S3 to S5** for *r*_*D*_, *r*_*E*_ and *r*_*I*_ of model simulations from all DBS frequencies.

To characterize the balance of excitation and inhibition in network models, we define the “inhibition strength ratio” (*ρ*_*inh*_) as the ratio of inhibitory to excitatory effective inputs. For a certain neural group “*m*” in the network, we define

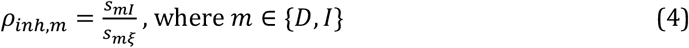

A network model that stresses the recurrent connections among excitatory nuclei is a “Hebbian network” ^24,42,55^. The effect from inhibitory nuclei is negligible in a Hebbian network ^42^, or incorporated in the background noise ^55^. Thus, we denote the network with small *ρ*_*inh*,*D*_ (i.e., dominant excitatory effects) as a Hebbian network (**Supplementary Table S6**). In the networks with relatively large *ρ*_*inh*,*D*_ (*ρ*_*inh*,*D*_ > 0.5), the inhibitory effective input (*s*_*DI*_) is an essential component of the network and is comparable with its excitatory counterpart. A network model with sufficient inhibitory effects in balancing excitatory effects is known as “Balanced Amplification network” ^24^, and we consider the network with *ρ*_*inh*,*D*_ > 0.5 (i.e., essential inhibitory effects) as a Balanced Amplification network. The network with the optimal model parameters (**Fig. 2**) is the typical Balanced Amplification network (*ρ*_*inh*,*D*_ = 0.91), in which the excitatory and inhibitory effects are equally strong. This optimal network minimizes the model fitting error (ER = 7.6%, **Supplementary Table S5**). To this end, we abbreviate “Balanced Amplification network” as “BA network” in text.

### Analysis of mechanism of Vim-network in response to DBS

The Vim-network mechanism is analyzed with the rate network model together with a further spiking network model (**Methods**) that we developed in this work. The spiking network model is a classical and ongoing approach for fitting the neural activity patterns in experimental data based on microscopic-level membrane potential dynamics ^22,40^; the commonly in-silico implemented Izhikevich spiking network model is a typical example ^22^. Based on the rate model parameters, we fitted the firing rate experimental data (**Fig. 2**) with an Izhikevich spiking network model (**Methods**) to explore the dynamics in clinical Vim-DBS data.

Similar to the rate model, the Izhikevich spiking network model was on three groups of neurons: (1) 20 Vim neurons (excitatory) directly receiving DBS; (2) 100 excitatory neurons from the cerebellum; (3) 40 inhibitory neurons from TRN. Compared with the rate model, in the spiking model, we incorporated more physiological details of the DBS effects and synaptic connections (**Methods**). Besides the DBS activation of the synapses afferent into the Vim neurons (**Fig. 1**), we incorporated the DBS effect of the axons efferent from the Vim neurons to the other neurons, and all synapses in the spiking network were characterized by the Tsodyks & Markram model ^36^ (**Methods**). The spiking model parameters were consistent with previous works ^22,56^, and were tuned so that for each group of neurons, the baseline firing rate (with DBS-OFF) was consistent with our rate network model (**Methods** and **Supplementary Table S8**). All spiking model simulations were conducted with the sampling resolution of 0.1 ms (same as the rate model). We simulated the spiking model with different frequencies (5∼200 Hz) of Vim-DBS, and obtained the instantaneous firing rate of the Vim neurons with a time histogram, which was computed with the optimized Gaussian kernel (same as the rate model, see **Supplementary Table S1**) on 20 simulated spike trains. We investigated two network mechanisms – Hebbian and Balanced Amplification – with the spiking network model; the two network mechanisms were characterized with the corresponding connectivity strength matrix W (Equation (1) and **Supplementary Table S4**), which is the result of our rate network model. To compare the two mechanisms, in the simulations, all the model parameters are the same except for the connectivity strength matrix W. We schematized the spiking network model mechanisms (**Fig. 3A**), and showed their impacts on the Vim firing rate (**Fig. 3C**); the spiking network model was compared with the rate network model (**Fig. 3B and D**). Note that in the spiking network, a cerebellar neuron is not connected with a TRN neuron ^37,39^ (**Fig. 3A, Methods**). In the rate network, there are recurrent connections among *E* and *I* groups of neurons, because *E* group is a more abstract structure incorporating cortical neurons (**Fig. 3B**).

**Fig. 3.**
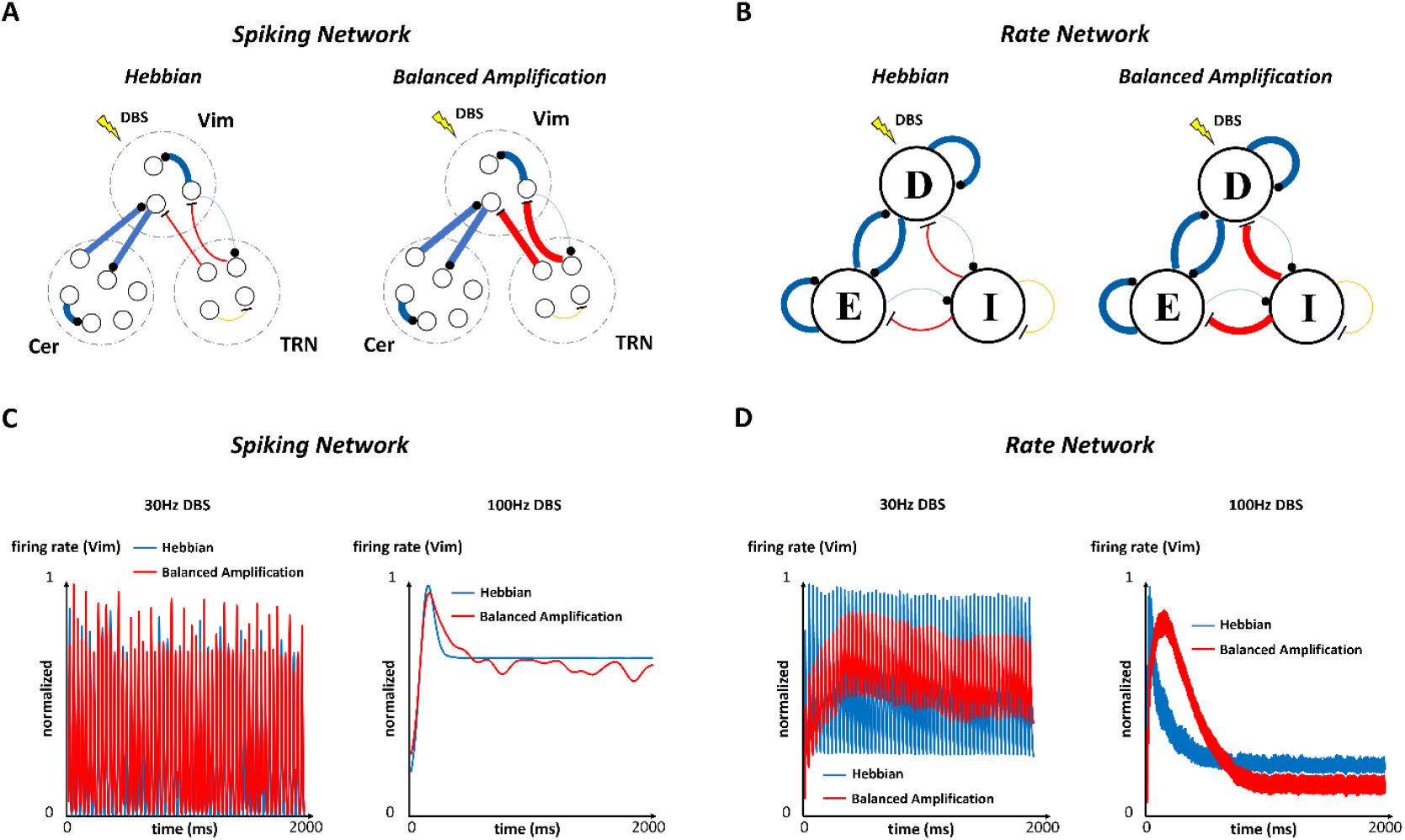
Mechanism analysis of Vim-network impacted by DBS. (A – B) Schematic representation of the network mechanisms. The synaptic connection with a dot (respectively, a bar) represents an excitatory (respectively, inhibitory) synapse. The boldness of the connection represents the effective input, which is the multiplication of connectivity strength and firing rate (Equation (3)). In the Hebbian mechanism, the ratio of inhibitory to excitatory effective inputs is low; such ratio is balanced (close to 1) in the Balanced Amplification mechanism. (A) Mechanisms of the spiking neuron network. The spiking network model consists of 3 groups of neurons: excitatory ventral intermediate nucleus (Vim) neurons directly receiving DBS, excitatory neurons in the cerebellum (Cer), and inhibitory neurons in the thalamic reticular nucleus (TRN). (B) Mechanisms of the rate network. The firing rate network model consists of 3 neural groups: “D” represents the Vim neurons directly receiving DBS, “E” represents the external excitatory nuclei, and “I” represents the external inhibitory nuclei. (C – D) Model simulations with the network mechanisms. We simulate the network models and present the results of Vim, receiving Vim-DBS of a typical low-frequency (3 Hz) and high-frequency (100Hz), respectively. We present the results from two network mechanisms: Hebbian and Balanced Amplification, corresponding to the mechanism schematics in (A) and (B). In each plot, the results are normalized to the maximum rate across the two mechanisms. (C) The spiking neuron network model simulations. (D) The firing rate network model simulations.

The spiking network model qualitatively represented the underlying mechanisms and the results of the rate network model (**Fig. 3C and D**). See **Supplementary Figs. S7 and S8** for the spiking model fits to data from different frequencies (5∼200 Hz) of Vim-DBS. In the spiking network, we found that the model fits with Hebbian and Balanced Amplification mechanisms were similar during 30-Hz DBS (a typical low-frequency DBS), but distinct during 100-Hz DBS (a typical high-frequency DBS) (**Fig. 3C**); consistent comparisons were observed in the rate network (**Fig. 3D**). The results from rate network model demonstrated that during low-frequency (≤50 Hz) DBS, the fitting accuracy with the Hebbian network is comparable to the BA network; this may implicate that the inhibitory neurons are not sufficiently engaged during weak inputs (low-frequency DBS) ^34^. However, during high-frequency (≥100 Hz) DBS, the Hebbian network deviates greatly from the BA network and experimental data (**Supplementary Table S7, Fig. 2 and Supplementary Fig. S6**). These results implicated that the Balanced Amplification mechanism (strong inhibition-stabilized effect) could be more evident during strong external inputs, e.g., a high-frequency of DBS pulses. This was consistent with the previous result on the ferret V1 neurons receiving various strengths of visual stimuli; it was hypothesized that the inhibition-stabilized effect dominated the network when the external visual stimuli were strong ^34^ .

During 100-Hz DBS, in the clinical data, we observed an initial large transient response lasting ∼600 ms and such transient response is accurately reproduced by the optimal BA rate network (**Fig. 2**). The existence and depression of the transient response can be explained by short-term synaptic plasticity (STP) ^18,12^. However, STP can’t explain the difference in the transient response between Hebbian and BA networks (**Fig. 3C and D**), where the same STP model and parameters are used in this work. In both rate and spiking network models, during 100-Hz DBS, we observed a shorter transient response in the Hebbian network than the BA network (**Fig. 3C and D**). A Hebbian network (small *ρ*_*inh*,*D*_, Equation (4)) stresses the recurrent excitation, which is less regulated by feedback inhibition ^24^. Thus, in the Hebbian network, the firing rate dynamics is less robust in response to strong external inputs like DBS. In a BA network (*ρ*_*inh*,*D*_ > 0.5), the strong external and recurrent excitation is stabilized by the strong feedback inhibition. In the optimal BA network, the inhibitory course has a longer time constant (τ_*i*_, table in **Fig. 2**) than excitatory effect (τ_*e*_, table in **Fig. 2**), and this may explain the longer initial transient response to 100-Hz DBS, compared with an excitation-dominant Hebbian network (**Fig. 3C and D**).

In the steady state firing rate in Vim clinical data, we observed some oscillatory dynamics during high-frequency DBS (**Fig. 2**). Such dynamics were observed in the spiking BA network, but almost completely missing in the spiking Hebbian network (**Fig. 3C**). These oscillatory dynamics can be explained by the Balanced Amplification mechanism stressing the critical role of inhibitory neurons, whose functions are hypothesized as follows. During 100-Hz DBS, due to the consecutive stimulation, Vim firing rate increased to a maximum at ∼200 ms, when the inhibitory neurons started to be sufficiently engaged and brought down the Vim firing rate after 200 ms. Then, the decreased Vim firing rate reduced the excitation to inhibitory neurons, whose firing rate decreased and thus disinhibited Vim neurons. So, Vim firing rate increased again, and this restored the engagement of inhibitory neurons. Such interplay between excitatory and inhibitory neurons formed the oscillatory activities in the steady-state response during 100-Hz Vim-DBS (**Fig. 3C** – model, and **Fig. 2** – clinical data).

The role of inhibitory neurons is further observed in the single-unit membrane potential data recorded in Vim neurons receiving high-frequency DBS (**Fig. 4**). We observed significant hyperpolarizing effects of inhibition, during both 100-Hz and 200-Hz DBS (**Fig. 4**). These evoked inhibitory activities were seldom observed during low-frequency (≤50Hz) DBS ^33^. Such a biomarker of inhibition starts to take effect after about 50 DBS pulses, which may indicate the time when the firing rate reaches the maximum and starts to decrease (**Fig. 2**).

**Fig. 4.**
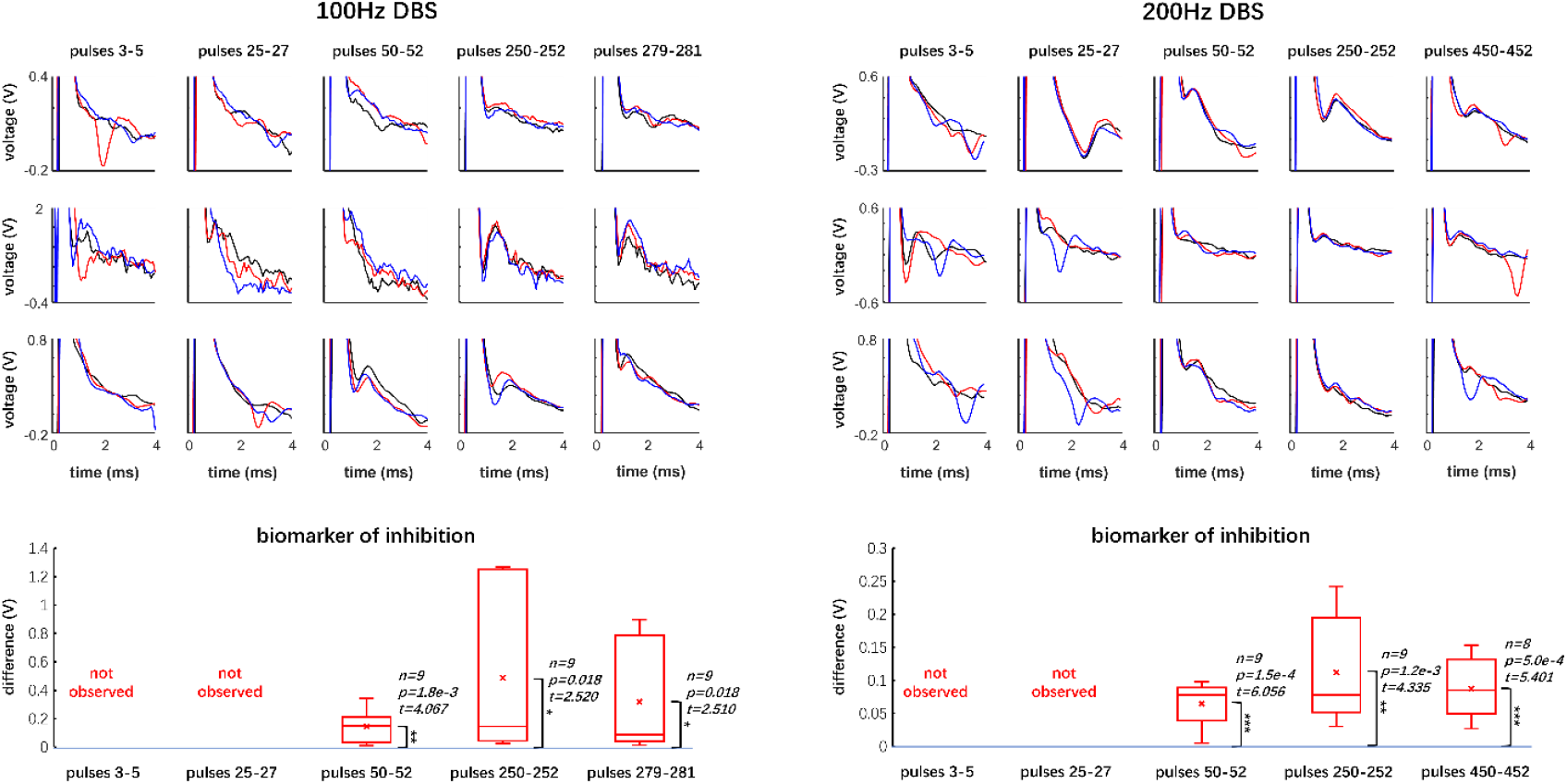
Biomarker of inhibition observed in single-unit membrane potential data. Single-unit recordings of the membrane potential of Vim neurons receiving DBS of high frequencies (100Hz and 200Hz). The recording corresponding to DBS pulse #i is from the inter-DBS-pulse-interval between the i^th^ and (i+1)^th^ DBS pulses. We present the recordings in the initial 4ms of each inter-DBS-pulse-interval. For each DBS frequency, we obtained recordings from 3 neurons, corresponding to each of the 3 rows. In each set of DBS pulses (e.g., pulses 3-5), the recordings are denoted with 3 colors: black, red and blue, corresponding to the ascending indices of the DBS pulses. For example, in pulses 3-5 from 100Hz DBS data, “the black, red and blue traces” denote data from “pulse 3, 4, and 5”, respectively. In each recording, the initial 1ms of data is truncated because it denotes the DBS artifact. The evident depolarizing feature, e.g., the red trace of the 1^st^ neuron (row) receiving pulses 3-5 of 100Hz DBS, represents the spiking activity. The evident hyperpolarizing feature, e.g., the 3 traces of the 2^nd^ neuron (row) receiving pulses 250-252 of 100Hz DBS, represents the inhibitory effects. The biomarker of inhibition is characterized by the difference between the peak of hyperpolarization and a preceding “valley” representing the initialization of inhibition (**Supplementary Table S11**). Prior to DBS pulse 50, the biomarker of inhibition is generally not observed. The biomarker is significantly observed in the shown data starting from DBS pulse 50. In the box-whisker plot, “x” marks the mean value of the biomarker. We use a one-sample one-tailed t-test to show that the biomarker of inhibition is significantly greater than 0. “n” represents the number of recordings used in the t-test. n=9 for all scenarios (3 pulses × 3 neurons), except for “200Hz DBS, pulses 450-452”, where the blue trace in the 3^rd^ row is excluded because the biomarker is hidden by the spiking (depolarizing) activity (**Supplementary Table S11**). One-sample one-tailed t-test: *p-value < 0.05; ** p-value < 0.01; *** p-value < 0.001

### Route optimization method and evolution of mechanisms

The rate model parameters (Equation (1)) were obtained from an optimization method that we developed (**Methods**). The network mechanism evolves from Hebbian to Balanced Amplification during the optimization process (**Fig. 1**). Our method, referred to as route optimization, effectively navigates the initial parameters in a route towards the globally optimal solution in a relatively large parameter space (the undetermined parameter set Φ in Equation (1) is relatively high dimensional with dim = 10). During the optimization process, we reduce the mean squared errors (MSE) between the experimental instantaneous firing rate and model predicted firing rates of the Vim neurons directly receiving DBS (group “D” in **Fig. 1**) (**Supplementary Notes 1**). Our aim was to find the optimal parameters that consistently fit the model to the instantaneous firing rate of Vim neurons receiving DBS across varying frequencies (5 to 200 Hz), where high-frequency (100 and 200 Hz) DBS are more stressed because they are more clinically effective ^8,9^.

We design two main stages of optimization, namely, Global Stage and Refining Stage in the route optimization method (**Fig. 5A**). Compared to Global Stage, Refining Stage stresses high-frequency DBS data to a higher extent (**Methods**). The Global Stage explores the parameter space and leads the route towards a “low error area”, which is exploited by the Refining Stage to locate the optimal parameters (**Fig. 5A**). Within each stage, the algorithm iterates between two objective functions, referred to as “Pushing Function” and “Stabilizing Function” (**Fig. 5B**), to estimate model parameters across different DBS frequencies. The objective functions are defined by the weighted sum of the MSE of individual DBS frequencies (**Supplementary Notes 1**). We use the term “Focused Feature” to represent the higher weights of the MSE of high-frequency (100-Hz and 200-Hz) DBS data (**Fig. 5B, Supplementary Notes 1**). Compared to Focused Feature, the term “Stabilized Feature” represents more balanced weights of the MSE of both low and high-frequency DBS (**Fig. 5B, Supplementary Notes 1**). Focused Feature is important in increasing the fitting accuracy during high-frequency DBS, which is underfitted in our previous study ^13^. Stabilized Feature is important to keep the balance of the fitting accuracy across all DBS frequencies. Pushing Function emphasizes Focused Feature, whereas Stabilized Feature is stressed in Stabilizing Function. Within each iteration in either Global Stage or Refining Stage, the two objective functions were sequentially executed: from the Pushing Function, the output parameters (Φ_*F*_, “F” represents Focused Feature) are the inputs to the Stabilizing Function (Φ_*S*_, “S” represents Stabilized Feature) (**Fig. 5B**). The output parameters of the Stabilizing Function will become the inputs to the Pushing Function in the next iteration (**Fig. 5B**). Pushing Function (on Focused Feature) “pushes” the model parameters away from the local solution towards the parameter range better representing Focused Feature, then Stabilizing Function (on Stabilized Feature) “stabilizes” the model parameters by fitting consistently across the whole data (i.e., Stabilized Feature) with high accuracy (**Fig. 5A, Methods**). Essentially, the pushing-and-stabilizing strategy keeps updating and improving the prior knowledge of the initial model parameters for each iteration of fits. This sequential process continues until the total fitting error (ER) (Equation (2)) was sufficiently small, and satisfies an exit rule (**Fig. 5B**, see Equation (22) in **Supplementary Notes 1** for details). The optimal parameters (Φ_*optimal*_) are the output of the route optimization method. See **Fig. 5A** and **Methods** for more details of the iterations during the optimization process.

**Fig. 5.**
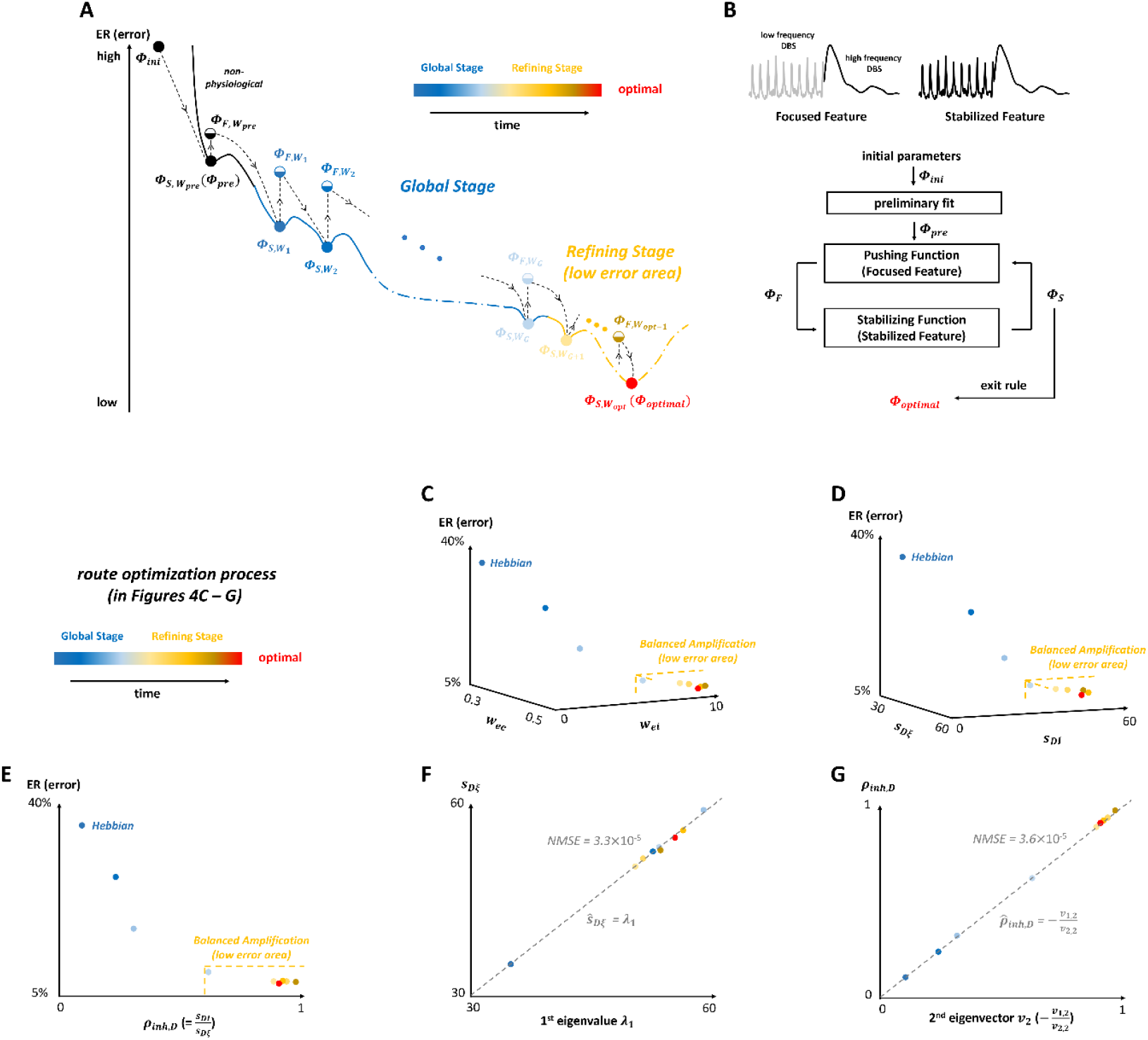
Route optimization method and evolution of mechanisms. (A) Schematic illustration of the route optimization method. From a set of initial parameters (Φ_ini_), we do a preliminary model fit and obtain the parameters Φ_pre_ . Φ_ini_ and Φ_pre_ that are in the “non-physiological” parameter range, meaning that the fitting errors (ER, Equation (2)) with these parameters are very high (**Methods**). The main stages of the optimization process are Global Stage and Refining Stage. Compared to Global Stage, Refining Stage stresses high-frequency DBS data to a higher extent. During the optimization process, Φ_F_ represents the model parameter set fitted to the Focused Feature (stressing high-frequency DBS data), and Φ_S_ is fitted to the Stabilized Feature (balancing low and high-frequency DBS data). W_i_ is the connectivity strength matrix (Equation (1)) in 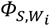, which evolves to 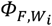 in the next model fit. 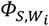 is represented by a fully filled circle, and 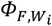 is represented by a half-filled circle of the same color as 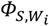; this means that part of the parameters (specifically, W_i_) are the same between 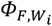 and 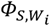 (**Methods**). (B) Route optimization flow chart. Pushing Function emphasizes the Focused Feature, and Stabilizing Function emphasizes the Stabilized Feature. The sequential iterations terminate after an exit rule (i.e., small fitting error, see Equation (22) in **Supplementary Notes 1**) is satisfied, and the final output is the optimal parameter set Φ_optimal_ that minimizes ER (error). (C – E) The model parameter set associated with the leftmost Hebbian network is 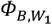 (in (A), **Supplementary Table S4**), which is the start of the Global Stage. (C) Evolution of the dominant connectivity strengths (w_ee_ and w_ei_, Equation (1), **Supplementary Table S4**). (D) Evolution of the effective synaptic inputs (s_Dξ_ and s_DI_, Equation (3), **Supplementary Table S6**). (E) Evolution of the inhibition strength ratio (ρ_inh,D_, Equation (4), **Supplementary Table S6**). (F) 1^st^ eigenvalue (λ_1_) of the effective input matrix (Equation (3)) and the excitatory effective input (s_Dξ_). “NMSE” is the normalized mean squared error. ŝ _Dξ =_ λ_1_ is a simple linear estimate. (G) 2^nd^ eigenvector (**v**_**2**_) of the effective input matrix (Equation (3)) and the inhibition strength ratio (ρ_inh,D_). 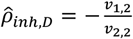 is a simple linear estimate.

As the route optimization evolved, the total fitting error ER (Equation (2)) decreased, and both *w*_*ei*_ and *s*_*DI*_ evidently increased, representing an increasing inhibitory effect (**Fig. 5C and D**). *w*_*ei*_ and *w*_*ee*_ are the dominant connectivity strengths (**Fig. 2, Supplementary Table S5**). As shown in **Fig. 5E**, ER decreased fast as *ρ*_*inh*,*D*_ increased from ∼0.1 to ∼0.6, then ER stabilized and reached the minimum when *ρ*_*inh*,*D*_∼0.9 (**Supplementary Tables S5 and S6**). Initially, the very low *ρ*_*inh*,*D*_ (∼0.1) represented a Hebbian network, where the excitatory effective input (*s*_*D*ξ_) is dominant and the inhibitory effective input (*s*_*DI*_) is negligible (**Fig. 5D and E**). As the optimization proceeds, the network evolved to the Balanced Amplification mechanism (*ρ*_*inh*,*D*_ > 0.5, **Fig. 5E**), with the optimal BA network (*ρ*_*inh*,*D*_= 0.91) representing equally strong excitation and inhibition. **Supplementary Video S1** illustrates the evolution of mechanisms during the route optimization process as shown in **Fig. 5E** (specified in **Supplementary Table S6**).

We implemented the inhibition strength ratio (*ρ*_*inh*,*D*_) to quantify the network mechanisms; small *ρ*_*inh*,*D*_ represents a Hebbian network, whereas large *ρ*_*inh*,*D*_ represents a BA network. *ρ*_*inh*,*D*_ was determined by the network excitatory and inhibitory effective inputs (*s*_*D*ξ_ and *s*_*DI*_) into the Vim neurons directly receiving DBS. Now the question is: Can *ρ*_*inh*,*D*_ represent the network features identified by the effective input matrix *S* (Equation (3)), which characterizes all nuclei in the modeled neural network? The answer is yes, substantiated by the following analysis in the perspective of the eigenvalues and eigenvectors of *S*.

The effective input matrix *S* consists of two eigenpairs: (λ_1_, ***v***_**1**_) and (λ_2_, ***v***_**2**_), where λ_*j*_ is the j^th^ eigenvalue, and the associated j^th^ eigenvector is 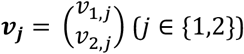. We analyzed the evolution of eigenpairs during the route optimization process (**Methods, Supplementary Table S6**). We found that the 1^st^ eigenpair (λ_1_, ***v***_**1**_) characterized *s*_*D*ξ_, i.e., the excitatory effective input into the Vim neurons directly receiving DBS; we deduced that *s*_*D*ξ_ ≈ λ_1_ (**Methods, Fig. 5G**). The 2^nd^ eigenpair (λ_2_, ***v***_**2**_) characterized *ρ*_*inh*,*D*_, i.e., the ratio of inhibitory to excitatory effective inputs to the Vim neurons directly receiving DBS (Equation (4)); we deduced that 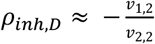 (**Methods, Fig. 5G**). We quantified these relationships by performing linear regression fits. The regression line of *s*_*D*ξ_ ∼ λ_1_ was

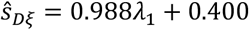

where the 95% confidence interval of the slope and vertical intercept was, respectively [0.962, 1.015] and [-0.977, 1.777]. Thus, this regression line very closely approximated ŝ_*D*ξ_ = λ_1_ (**Fig. 5G**, gray dashed line). Similarly, the linear regression fit result of 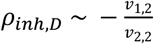 was

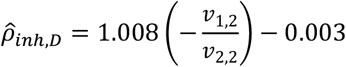

where the 95% confidence interval of the slope and vertical intercept was, respectively [1.002, 1.014] and [-0.007, 0.002]. This result was almost identical to 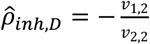 (**Fig. 5G**, gray dashed line). For both simplified linear fits 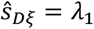 and 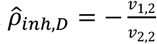, the NMSE was smaller than 10^-4^ (**Fig. 5F and G**). This demonstrated that the full variation of *s*_*D*ξ_ (respectively, *ρ*_*inh*,*D*_) was captured by λ_1_ (respectively, 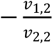) in all networks, with either Hebbian (small *ρ*_*inh*,*D*_) or Balanced Amplification (large *ρ*_*inh*,*D*_) features.

The analysis with the eigenvectors and linear regressions demonstrated that the network features of *S* can be characterized by *s*_*D*ξ_ and *ρ*_*inh*,*D*_. In **Fig. 5G**, we saw that as the route optimization evolved, the dynamics of *s*_*D*ξ_ was random, except for the initial smaller value. Despite such randomness in *s*_*D*ξ_, its dynamics could always be perfectly captured by the 1^st^ eigenvalue λ_1_ (ŝ_*D*ξ_ = λ_1_, **Fig. 5G**). As the route optimization evolved, *ρ*_*inh*,*D*_ first increased then stabilized around the optimal value at ∼0.9; the dynamics of *ρ*_*inh*,*D*_ could always be perfectly captured by the 2^nd^ eigenvector ***v***_**2**_(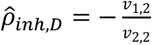, **Fig. 5G**).

## Discussion

### The route optimization method is automatic and accurate

We developed a model parameter optimization method that constructs a route leading to the globally optimal solution. Our route optimization method is automatic, in the sense that there is no hand-tuning of the model parameter set *Φ* (Equation (1)) throughout the optimization process. Automatic parameter optimization is more accurate and efficient than the parameter hand-tuning, which is the most often implemented approach in analyzing network models or fitting data ^57,26,58^. Although easy to explore, the parameter hand-tuning is often laborious and complicated to be implemented to find appropriate results ^59^, and depends heavily on prior knowledge which may not be optimal for the current data ^26,58^. With an effective automatic optimization method, we can efficiently find the optimal model parameters that accurately fit an experimental dataset, and these optimal parameters probably characterize the physiological underpinnings. During our optimization, we observed that the high fitting accuracy requires the large value of the variable *w*_*ei*_ (Equation (1), table in **Fig. 2**), which represents the synaptic connectivity strength from inhibitory neurons to excitatory neurons. *w*_*ei*_ is the critical quantity representing the inhibition-stabilized effect. By validating sufficient rounds of simulations, we confirmed that the model fitting accuracy was low if *w*_*ei*_ was not large enough (data not shown). These observations from optimization processes further imply that inhibitory effects are essential in fitting Vim-DBS data and further characterizing the Vim-network.

### Clinical evidences of Balanced Amplification mechanism

The Balanced Amplification mechanism stressing the critical role of inhibitory neurons has been validated in experimental works of different brain circuits ^60,61,62^. In-vitro brain-slice experiments showed that in response to external stimuli, in general, the depression effect can be more evident for excitatory (glutamatergic) synapses than inhibitory (GABAergic) synapses, and E/I balance shifted towards inhibition as the stimulation frequency increases ^61,62,63,64,65^. In the thalamic circuits, the main inhibitory input to thalamic relay nuclei is from thalamic reticular nuclei (TRN) ^66^. With in-vitro slice recordings, Campbell et al. (2020) ^60^ demonstrated that the firing rate of thalamic relay nuclei significantly decreases as the optogenetic stimulation frequency increases. The significant firing rate reduction implied that E/I was shifted towards inhibition during higher frequency of stimulation, and the possible mechanism was further demonstrated by analyzing the recorded inhibitory post-synaptic potential (IPSP) of the thalamic relay nuclei ^60^. The levels of hyperpolarization (i.e., IPSP amplitude) during high-frequency stimulation is significantly larger than during low-frequency stimulation ^60^. This showed that the inhibitory effect from TRN is resilient and significantly increased under high-frequency stimulation ^60,67^. These experimental results are consistent with the prediction from our model: in the simulations of both rate and spiking network models, we observed that the Balanced Amplification mechanism is more evident during high-frequency DBS than low-frequency DBS (**Fig. 3**). Clinical observations demonstrated that thalamic-DBS with low-frequency (≤50 Hz) can exacerbate essential tremor ^68,69^. The appearance of tremor – in the disease state and low-frequency DBS – may indicate an underlying lack of Balanced Amplification mechanism (inhibition stabilization effects) of otherwise suppressed excitatory oscillations – these oscillations manifest as tremors in the muscles. Thus, a possible therapeutic mechanism of high-frequency Vim-DBS is to restore the Balanced Amplification that could have perhaps been lost due to the disease state or low-frequency DBS.

### Potential generalizability of mechanism analysis to other circuits

We wanted to investigate the compatibility of our modeling strategy with different types of neuromodulations besides DBS. Another study observed the experimental single-unit recordings in ferret primary visual cortex (V1) neurons receiving visual stimuli with various stimulation strengths (e.g., different contrast or stimulus size) ^34^. As the visual stimulation strength increased, the steady-state firing rate of ferret V1 neurons increased supralinearly when the stimuli were weak, and started to decrease when the stimuli became strong (see Figs. 4 and S10 in Rubin et al. (2015) ^34^). Such firing rate dynamics in response to various strengths of visual stimuli was modeled as a “stabilized-supralinear network”: as external input strength increases, the network transits from dominantly externally driven (supralinear increase in firing rate) to dominantly network driven, with the network inhibition stabilizing effect becomes increasingly dominant ^34^. This stabilized-supralinear network model was developed and implemented to qualitatively represent the ferret V1 neuron experimental recordings in response to various strengths of visual stimuli ^34^. In Vim-DBS (**Supplementary Fig. S10A**), during low-frequency DBS (5 to 50 Hz), the steady-state firing rate of the group *D* neurons (Vim neurons directly receiving DBS) increased as the DBS frequency increased; the steady-state firing rate decreased during high-frequency DBS (100 and 200 Hz). Such dynamics of experimental steady-state firing rate in response to various frequencies of DBS was accurately reproduced by our rate network model (**Supplementary Fig. S10A and C**). Besides, human Vim and ferret V1 neurons reacted similarly to the external drive (DBS and visual stimulation, respectively); as the strength of the external drive increased (increasing DBS frequency or visual stimulation strength), the steady-state firing rate first increased, then fast decreased. This implies that our modeling framework – including the rate network model, optimization method and mechanism analysis strategy – can be potentially generalizable to studying visual cortex and various brain circuits, receiving different types of external stimuli.

### Limitation of study

The DBS experimental data implemented in this work were recorded in a relatively short time scale (≤10 s) for each stimulation frequency (5∼200 Hz) (**Methods**). Within these short windows of recorded activity immediately following each DBS pulse, we divide the neuronal temporal response profile into an early-transient response and a latter steady-state response (**Fig. 2**) within the context of clinical DBS for essential tremor. Since tremor symptoms often respond to stimulation on the scale of seconds, the relatively short timescales of our experimental data are nonetheless likely sufficient for initializing optimal DBS settings in clinics ^70,71^. However, in considering the effects of stimulation at a longer time scale (≥ minutes), the stimulated neurons may exhibit features of long term synaptic plasticity (LTP), which is the long-term change in the synaptic connectivity or morphology ^72,73^. LTP is widely observed in cortical neural circuits pertinent to memory ^72^, learning ^73^ and neuromodulations ^73^, etc. The modeling of the possible LTP in long-duration DBS is part of our future work.

Our model extracts the main architecture of the Vim-circuitry, i.e., the Vim neural group and its recurrent connections with external excitatory and inhibitory neural groups. We incorporated all the external excitatory (resp., inhibitory) neurons into one group. Such structure blurred the details of these excitatory (resp., inhibitory) neurons. The connections among the neural groups were modeled by static synapses, i.e., the synaptic connectivity strength – e.g., *w*_*ei*_ (Equation (1)) – is not time-varying. We incorporated synaptic plasticity (characterized by the Tsodyks & Markram model ^36^) in the DBS-induced inputs into the Vim neural group, but synaptic plasticity was not considered within the modeled network. The purpose for such abstraction is to reduce the dimensionality and complexity of the rate model. We found that the robustness of model simulation decreases if incorporating more variables of synaptic connectivity into the model (data not shown). Thus, in the rate model, due to lack of synaptic plasticity within the modeled network, we didn’t explicitly incorporate the DBS effect of the axons efferent from the Vim neurons towards external neural groups. The modeling of detailed synapses – including the DBS effect of Vim efferent axons – was incorporated in the spiking network model (**Fig. 3** and **Methods**) that we developed.

### Conclusions and future work

We developed a firing rate network model of the Vim-circuitry impacted by Vim-DBS, and a parameter route optimization method that automatically and effectively found the globally optimal model parameters fitted accurately and consistently across experimental data from various DBS frequencies (5∼200 Hz). Inferred from these optimal model parameters, we detected the Balanced Amplification mechanism characterizing the strong inhibition stabilization effects underlying the Vim-circuitry. Our modeling, parameter optimization and mechanism analysis strategies are potentially generalizable to studying various brain circuits, receiving different types of neuromodulations besides DBS. The current work could be extended to construct detailed macroscopic thalamocortical network models on quantifying different neurological disease mechanisms (**Supplementary Notes 2**), and these future models can be potentially implemented in developing model-based closed-loop DBS control systems ^41^ for automatic disease treatments.

## Methods

### Human experimental data

We used the same human experimental single-unit recordings which were previously published in Milosevic et al. (2021) ^12^. Thus, the commitment to ethics policies have already been fulfilled ^12^. All human experiments conformed to the guidelines set by the Tri-Council Policy on Ethical Conduct for Research Involving Humans and were approved by the University Health Network Research Ethics Board ^12^. Moreover, each patient provided written informed consent prior to taking part in the studies ^12^.

The human experimental data protocols and offline analysis methods were from Milosevic et al. (2021) ^12^. Microelectrodes were used to both deliver DBS and perform single-unit recordings. DBS was delivered using 100 µA and symmetric 0.3ms biphasic pulses (150µs cathodal followed by 150 µs anodal) ^12^. In this work, we used the single-unit recordings of the neurons in the thalamic ventral intermediate nucleus (Vim) of essential tremor patients, during various DBS frequencies (5 to 200 Hz) in Vim. The single-unit recordings during {5, 10, 20, 30, 50, 100, and 200 Hz} Vim-DBS are of length {10, 5, 3, 2, 1, 5, and 2 s}, respectively; for each frequency of DBS, we did 5 to 8 recordings in different patients (total number of patients = 19). To obtain spikes from the single-unit recordings, we did offline analysis and spike template matching. For each single-unit recording, all the narrow stimulus artifacts were removed (0.5 ms from the onset of a DBS pulse). Then the recordings were high pass filtered (≥300 Hz) to better isolate the spikes, which were identified by the template matching using a principal component analysis method in Spike2 (Cambridge Electronic Design, UK).

### Time histogram with the optimized Gaussian kernel

After offline processing of the Vim single-unit recordings, we obtained 5 to 8 spike trains for each frequency of Vim-DBS (5, 10, 20, 30, 50, 100, and 200 Hz), and computed the corresponding instantaneous firing rate with a time histogram method, which is a common practice in processing spiking data. However, people usually subjectively choose the kernel width for the time histogram, and ignore the underlying mechanisms generating the spikes ^52^. In our work, we computed the instantaneous firing rate by convolving the spike trains with the optimized Gaussian kernel, which was obtained by the method in Shimazaki et al. (2007) ^52^ and Shimazaki et al. (2010) ^53^. The optimized Gaussian kernel characterized the true Poisson process underlying the experimental spiking data ^52,53^. Specifically, the optimized kernel minimized the mean integrated square error (MISE) from the true inhomogeneous Poisson point process underlying the experimental spike trains ^52,53^. A time histogram kernel with small MISE could capture the abrupt firing rate fluctuations, while reducing the excessive overfitting ^52,53^. From the spike trains recorded during each frequency of DBS, we computed the instantaneous firing rate with the optimized Gaussian kernel; see **Supplementary Table S1** for the optimized kernel of the data from each DBS frequency. The instantaneous firing rate with the optimized Gaussian kernel grasped the transient fluctuations very well, while the overfitting was prevented, in both 5-Hz and 200-Hz DBS data (**Supplementary Fig. S1**). Compared with the optimized Gaussian kernel, a larger kernel caused the instantaneous firing rate to deviate from the transient fluctuations, and a smaller kernel made the instantaneous firing rate exhibit excessive overfitting (**Supplementary Fig. S1**).

### The Tsodyks & Markram model of short-term synaptic plasticity (STP)

The external input into the Vim-network was the DBS-induced post-synaptic current (*I*_*DBS*_, **Fig. 1**, Equation (1)), which was formulated with the Tsodyks & Markram (TM) model of short-term synaptic plasticity (STP) ^36^. Thus, the immediate impact of DBS pulses was modeled as inducing synaptic release, and such modeling method is consistent with previously established works ^12,13,74^. For the Vim neurons directly receiving DBS, we modeled that each neuron receives inputs from 500 synapses, with 90% excitatory synapses (*N*_*exc*_ = 450) and 10% inhibitory synapses (*N*_*inh*_ = 50) (^12^, **Fig. 1, Supplementary Table S2**). We assumed that each DBS pulse generates one spike in each of these synapses simultaneously ^12,13^. These DBS-evoked spikes were filtered by the TM model, generating the post-synaptic current, *I*_*DBS*_, that was obtained by a linear combination of post-synaptic excitatory (*I*_*exc*_) and inhibitory (*I*_*inh*_) currents as follows:

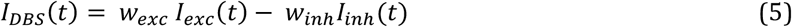

where *w*_*exc*_ and *w*_*inh*_ denote the scaling weights of the excitatory and inhibitory currents, respectively (**Supplementary Table S2**). *I*_*exc*_ (respectively, *I*_*inh*_) is the total post-synaptic current from all excitatory (respectively, inhibitory) synapses; each synapse (excitatory or inhibitory) was modeled by the TM model of short-term synaptic plasticity:

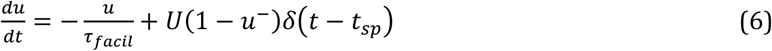

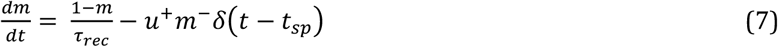

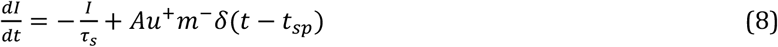

where *u* is a utilization parameter, indicating the fraction of neurotransmitters ready for release into the synaptic cleft (due to calcium ion flux in the pre-synaptic terminal). The variable *m* indicates the fraction of resources remaining available after the neurotransmitter depletion caused by neural spikes. We denote as *u*^−^ and *m*^−^ the corresponding variables just before the arrival of the spike; similarly, *u*^+^ and *m*^+^ refer to the moment just after the spike. The *δ*–function modeled the abrupt change upon the arrival of each pre-synaptic spike *t*_*sp*_; for example, at *t* = *t*_*sp*_ in Equation (6), *u* increases by *U*(1 − *u*^−^), and *δ*(*t* − *t*_*sp*_) = 0 when *t* ≠ *t*_*sp*_. If there is no pre-synaptic activity (spike), *u* exponentially decays to zero; this decay rate is the facilitation time constant, τ_*facil*_ (Equation (6)). In contrast to the increase of *u* upon the arrival of each pre-synaptic spike, *m* drops and then recovers to its steady state value (= 1); this recovery rate is given by the recovery time constant τ_*rec*_ (Equation (7)). The competition between the facilitation (τ_*facil*_) and recovery (τ_*rec*_) time constants determined the dynamics of the synapse. In the TM model, *U*, τ_*facil*_, and τ_*rec*_ were the parameters that determined the three types of the synapse: facilitation (“F”), pseudo-linear (“P”), and depression (“S”) (^12^, **Fig. 1, Supplementary Table S2**). In Equation (8), *I* is the post-synaptic current, *A* is the absolute response amplitude, and τ_*s*_ is the post-synaptic time constant (**Supplementary Table S2**). We obtained *I*_*exc*_ (respectively, *I*_*inh*_) by adding the post-synaptic currents from all excitatory (respectively, inhibitory) synapses. The TM model parameters in **Supplementary Table S2** were chosen based on the previous modeling works on specific experimental datasets ^12,75,76^.

### Route optimization method for model parameters

The route optimization starts with a set of initial parameters (*Φ*_*ini*_), which came from previously reported parameters in other studies and a one-step fit with our previous single-ensemble model of the Vim neural group receiving DBS (^13^, **Supplementary Notes 1**). In particular, the initial connectivity strength parameters ({*w*_*ee*,_*w*_*ie*,_*w*_*ei*_, *w*_*ii*_}) were all set to 1. The output parameters from each optimization objective function (Pushing Function or Stabilizing Function) were obtained with the MATLAB custom function “fminsearch”, which implemented the Nelder-Mead simplex method ^77^. Starting from *Φ*_*ini*_, we did a preliminary fit with Stabilizing Function, and the output parameters were *Φ*_*pre*_ (**Fig. 5A and B**). The NMSE of the model fits with *Φ*_*ini*_ and *Φ*_*pre*_ were very large (>40%, **Supplementary Table S5**); thus, the networks with these model parameters were considered to be in the “non-physiological” regime, meaning that they were very different from the experimental data (**Fig. 5A**). *Φ*_*pre*_ was also denoted as 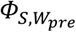 (**Fig. 5A**); the subscript “S” represents that it was the output of Stabilizing Function emphasizing the Stabilized Feature, and the subscript “*W*_*pre*_” stresses the strength of connectivity among neural groups in the network associated with *Φ*_*pre*_. Starting from 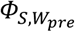, we performed a fit with Pushing Function, and obtained the output parameters 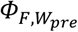 ; the subscript “F” represents that it was the output of Pushing Function that emphasizes Focused Feature (**Fig. 5A**). Note that in the fit for obtaining 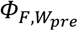 from 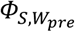, we fixed the connectivity strength *W*_*pre*_; and for 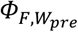, this feature (fixed *W*_*pre*_) is illustrated as a half-filled circle (**Fig. 5A**). During the route optimization process, the connectivity strength was left unchanged in the Pushing Function to obtain *Φ*_*F*_ from *Φ*_*S*_. Our purpose of this design was to increase the prediction robustness of the connectivity strength, which was the essential feature of a neural network. With the same connectivity strength *W*, the error of model fit with *Φ*_*F*,*W*_ was often larger than *Φ*_*S*,*W*_ (**Fig. 5A, Supplementary Table S5**) because *Φ*_*F*,*W*_ was obtained during fitting Focused Feature stressing part of the data (high-frequency DBS). However, from *Φ*_*S*,*W*_ to *Φ*_*F*,*W*_, the model parameters always moved away from the local minimum, and the chances were opened up to explore the global minimum satisfying our optimization objective (**Fig. 5A**). Thus, Pushing Function (on Focused Feature) “pushes” the model parameters away from the local solution towards the parameter range better representing Focused Feature, then Stabilizing Function (on Stabilized Feature) “stabilizes” the model parameters by fitting consistently across the whole data (i.e., Stabilized Feature) with high accuracy. Such sequential execution of Pushing Function and Stabilizing Function (**Fig. 5B**) is the critical idea of the route optimization method in effectively finding the globally optimal solution.

Starting from *Φ*_*pre*_, the 1^st^ sequential optimization iteration is 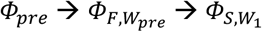 (**Fig. 5A and B**). The model fitting error (ER) of the network with parameters as the 1^st^ iteration output 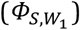 was 35.9%, which is far smaller than the networks in the non-physiological regime (**Fig. 5A, Supplementary Table S5**). Thus, we considered the network associated with 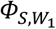 as physiologically meaningful, as well as each of the following network associated with a subsequent iteration output 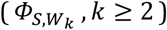; for these networks, ER decreases as the iteration number *k* increases (**Fig. 5A, Supplementary Table S5**). In the physiological regime, starting from 1^st^ iteration output 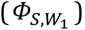, the route optimization proceeds as the sequential iteration number increases; the *k*^th^ iteration (*k* ≥ 2) is 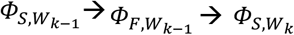 (**Fig. 5A**). The route optimization process in the physiological regime consists of two stages: Global Stage and Refining Stage (**Fig. 5A**). The difference between the two stages is that, Refining Stage emphasizes the high-frequency DBS data to a greater extent in the objective functions (**Supplementary Notes 1**). Compared to Global Stage, in Refining Stage, the weight of high-frequency DBS data was greater in both Pushing Function and Stabilizing Function (**Supplementary Notes 1**). The purpose of Global Stage is to navigate the model parameters towards the appropriate range for fitting the data across all DBS frequencies; Refining Stage refines the parameter range obtained by Global Stage to further reduce the fitting error, in particular of the more clinically effective high-frequency DBS data. The fitting error of the network associated with the final output of Global Stage (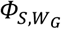, **Fig. 5A**) was already small (ER = 9.5%, NMSE (100 Hz) = 4.3%, NMSE (200 Hz) = 10.6%); however, utilizing the objective functions of Global Stage solely would cause ER to stop decreasing (data not shown). Thus, we switched to Refining Stage to fully explore and exploit the “low error area” (**Fig. 5A**) to minimize ER; after such local refinement, we finally found the desired globally optimal solution *Φ*_*optimal*_ (ER = 7.6%, NMSE (100 Hz) = 3.2%, NMSE (200 Hz) = 5.7%), which satisfied an exit rule (**Supplementary Notes 1**) to quit the optimization iterations (**Fig. 5B**). The optimal model fit the associated model parameters (*Φ*_*optimal*_) were shown in **Fig. 2**.

### Eigenpairs of the effective input matrix

We computed the eigenpairs – eigenvalues and the associated eigenvectors – of the effective input matrix *S* (Equation (3)); **Supplementary Table S6** showed the evolution of *S* and its eigenpairs during the route optimization process (as illustrated in **Fig. 5F and G**). Suppose that the *i*^*th*^ eigenpair of the effective input matrix *S* is (λ, 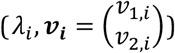), and we deduced the following equations:

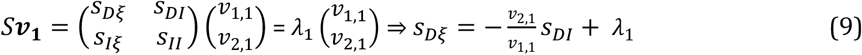

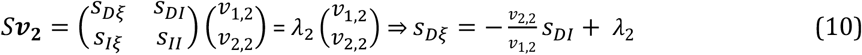

where λ_*i*_ (*i* = 1,2) is the *i*^*th*^ eigenvalue of *S*, and 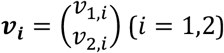 is the *i*^*th*^ eigenvector of *S* associated with λ_*i*_ .

During the route optimization process, we always observed that 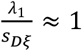 and 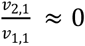 (**Supplementary Table S6**). This observation was consistent with Equation (9), which concretely showed that *s*_*D*ξ_ ≈ λ_1_ when 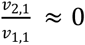. Thus, the excitatory effective input (*s*_*D*ξ_) could mostly be represented by the 1^st^ eigenvalue (λ^1^). When observing the 2^nd^ eigenpair (λ_2_, ***v***_**2**_) during the route optimization process, we found that 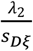 is always small; in fact, max 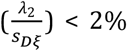 (**Supplementary Table S6**). Thus, in analyzing the inhibition strength ratio 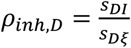 (Equation (4)), we deduced the following equation from Equation (10):

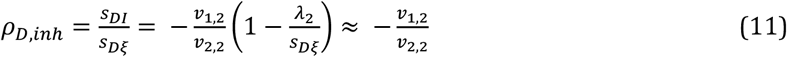

because max 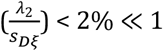 (**Supplementary Table S6**). Thus, the inhibition strength ratio could mostly be represented by the 2^nd^ eigenvector 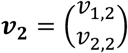.

### The Izhikevich spiking network model

We developed an Izhikevich spiking network model of the Vim-network, and compared its fitting results with our rate network model. The purpose was to validate our rate network model, and further discuss the network mechanism. The Izhikevich spiking network model was of 3 groups of neurons: (1) 20 Vim neurons (excitatory) directly receiving DBS; (2) 100 excitatory neurons from the cerebellum; (3) 40 inhibitory neurons from the thalamic reticular nuclei (TRN). This setup of the Izhikevich model is consistent with the previous spiking models of the thalamocortical network ^22,56^. The Izhikevich spiking network model equations are stated as follows,

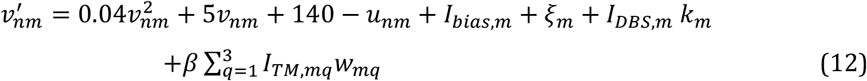

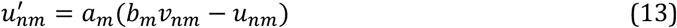

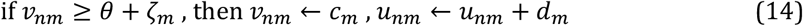

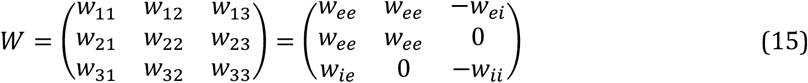

where m is the index of the neural group; m = 1, 2, 3 corresponds to the neural group Vim, cerebellum and TRN, respectively. Each group consists of *N*_*m*_ neurons, which are indexed by n = 1, 2, 3, …, *N*_*m*_ ; *N*_1_ = 20, *N*_2_ = 100, and *N*_3_ = 40. The simulation of the Izhikevich spiking network model was performed with the Neural Simulation Tool (NEST) platform ^78^.

In Equation (12), *v*_*nm*_ is the membrane potential of the n^th^ neuron in the m^th^ group, and *u*_*nm*_ is the corresponding membrane recovery variable. *I*_*bias*,*m*_ is the constant biased current injected to each neuron in the m^th^ group, and ξ_*m*_ is the Gaussian white noise (see **Supplementary Table S8** for its standard deviation). *I*_*DBS*,*m*_ is the DBS-induced post-synaptic current into the m^th^ group, with synapses characterized by the Tsodyks & Markram (TM) model ^36^; *k*_*m*_ is a scaling parameter. *I*_*DBS*,1_ represents the DBS activation of the synapses afferent into the Vim neurons (**Fig. 1**). To be consistent with the input to the Vim-network in our rate model, *I*_*DBS*,1_ was determined to be the same as the rate model input (*I*_*DBS*_) in Equation (1). *I*_*DBS*,2_ (respectively, *I*_*DBS*,3_) represents the DBS effect of the axons efferent from the Vim neurons into the cerebellar neurons (respectively, neurons in TRN). *I*_*DBS*,2_ and *I*_*DBS*,3_ were simulated by the NEST built-in function “tsodyks2_synapse”. We tuned the parameters in “tsodyks2_synapse” so that its simulation result is consistent with the Tsodyks & Markram synapses used in our rate model, across different DBS frequencies (5∼200 Hz) (**Supplementary Table S9, Supplementary Figs. S2 and S9**). We tuned *k*_1_ (the scaling of DBS effect of Vim afferent) so that the amplitude of the simulated DBS-evoked activities is consistent with our rate network model (**Fig. 2 and Supplementary Fig. S8**). The scaling parameters of DBS effect of Vim efferent (*k*_2_ and *k*_3_) are smaller than Vim afferent (*k*_1_), because there is energy loss during the transmission on the axons efferent from Vim ^79,80^. The ratio *k*_2_/*k*_3_ (i.e., the comparison of Vim efferent effects to cerebellum and TRN) is the same as the ratio of the corresponding connectivity strength (*w*_*ee*_/*w*_*ie*_, Equations (1) and (22)).

In Equation (12), *β* is a scaling parameter of the effect of synaptic connections within the network. *I*_*TM*,*mq*_ is the sum of TM-modeled post-synaptic currents induced by each spike of the group “q” neurons projecting to a group “m” neuron. *I*_*TM*,*mq*_ was simulated by the NEST built-in function “tsodyks2_synapse” with parameters specified in **Supplementary Table S9**. In the simulation of *I*_*TM*,*mq*_, we used the sparse synaptic connection, which is commonly used in spiking network models ^81,82^. In our simulation, there is no synaptic connection between a cerebellar neuron and a TRN neuron ^37,39^ (*w*_23_ = *w*_32_ = 0, Equation (15)). For the connected neurons, each neuron received synaptic inputs from 10% of Vim neurons (excitatory), 10% of cerebellar neurons (excitatory), and 25% of TRN neurons (inhibitory). The connection probability from an inhibitory neuron was set to be higher, because an inhibitory neuron is generally dense in axons, and can project axons to distal neurons ^83,39^. In the connectivity matrix W, *w*_*mq*_ represents the connectivity strength from the neural group “*q*” to group “*m*”, and the +/-sign denotes the excitatory/inhibitory effect (Equation (15)). The connectivity matrix W in the spiking model is the same as the rate model (Equation (1)). In particular, in the spiking model, the two mechanisms – Hebbian and Balanced Amplification – are characterized by the connectivity strengths shown in **Supplementary Table S4** (results of our rate model).

In Equation (13), *u*_*nm*_ is the membrane recovery variable, *a*_*m*_ is the time scale, and *b*_*m*_ is the sensitivity to the membrane potential *v*_*nm*_ . The neuron fires when *v*_*nm*_ reaches a fluctuating threshold *θ* + ζ_*m*_ (Equation (14)); *θ* = 30 mV is the mean firing threshold, and ζ_*m*_ is the Gaussian white noise (see **Supplementary Table S8** for its standard deviation). Immediately after the neuron fires, *v*_*nm*_ is reset to *c*_*m*_, and *u*_*nm*_ is reset to *u*_*nm*_ + *d*_*m*_ (Equation (14)). The Izhikevich model parameters {a, b, c, d} for the neurons in each group were shown in **Supplementary Table S8**; these parameters were chosen to be consistent with other works ^22,56^ . Other spiking model parameters were tuned so that for all neurons (Vim, cerebellum, or TRN), the simulated baseline firing rate (with DBS-OFF) was consistent with our rate network model.

### Statistical analyses

One-sample one-tailed t-test was performed to analyze the statistical significance of the biomarker of inhibitory effects observed in single-unit recordings, as shown in **Fig. 4**. Analysis of variance (ANOVA) test with F-statistic was performed to compare the length of initial transient responses between 100-Hz and 200-Hz DBS clinical data (**Supplementary Table S10**). A linear regression analysis was performed to generate the results related to **Fig. 5**. The regression methods include 95% confidence intervals of the slope and vertical intercept.

## Data availability

Human experimental data – Vim spike timings during different frequencies of Vim-DBS – have been deposited at https://github.com/nsbspl/rate_network_model.

## Code availability

The codes for generating results are openly accessible at https://github.com/nsbspl/rate_network_model.

## Acknowledgements

We acknowledge AmirAli Farokhniaee for helpful discussions in simulating the Izhikevich spiking network model. This work was supported by Milad Lankarany’s NSERC Discovery Grant (RGPIN-2020-05868), CIHR Project Grant and Brain Canada (Azrieli Foundation).

## Author contributions

Conceptualization, Y.T. and M.L.; Rate network model, Y.T. and M.L.; Optimization and data analysis, Y.T.; Result analysis, Y.T. and M.L.; Biological and clinical interpretations, Y.T., E.B., D.C., Z.P., L.M., and M.L.; Spiking network model, Y.T., D.C., L.M., and M.L.; Data collection, S.K.K., M.H., A.M.L., W.D.H., and L.M.; Writing – original draft, Y.T. and M.L.; Writing – review & editing, Y.T., E.B., D.C., Z.P., M.D.J., M.R.P., L.M., and M.L.; Funding acquisition, M.L.

## Materials & correspondence

Correspondence and requests for materials should be addressed to Milad Lankarany.

## Supplementary information

The supplementary materials – Supplementary Notes, supplementary figures, supplementary tables and a supplementary video – are included in a separate file.

