## Supplementary Information for "Uncovering network mechanism underlying thalamic deep brain stimulation"

#### Supplementary Notes 1 – route optimization

##### *Background and rationale*

The route optimization method finds the globally optimal solution in a relatively large parameter space. A globally optimal solution is important in fully fitting DBS experimental data, because the data features are distinct between low (≤50 Hz) and high (≥100 Hz) frequencies of DBS [1,2], and typical optimization methods focusing on a local solution [3,4,5,6,7] will be biased towards a single DBS data feature [2]. In this work, our optimization method incorporates two physiological features of the DBS data: (1) Focused Feature, stressing the Vim firing rate dynamics observed during high frequency Vim-DBS, which is more clinically effective for essential tremor [8,9,10]; and (2) Stabilized Feature, referring to the Vim firing rate dynamics across both low and high frequencies of Vim-DBS.

Local optimization methods may not be sufficient to fit both Focused Feature and Stabilized Feature in a relatively large parameter space (the undetermined parameter set $\Phi$ (Equation (1)) is relatively high dimensional with dim = 10). Local optimization methods might be effective in exploitation, i.e., exploiting a local parameter range for the suitable local solution, but are not sufficient in exploration, i.e., exploring the whole parameter space for a globally appropriate solution [3,5,6]. An optimization method for finding the globally optimal solution should be effective in both exploitation and exploration.

In the route optimization method, we navigate an exploiting optimizer in a route that effectively explores the whole parameter space for the globally optimal solution. For the exploiting optimizer, we used the Nelder-Mead simplex method in the MATLAB custom function "*fminsearch*" [6,7]. The Nelder-Mead method is one of the most accurate gradient-approximating optimization methods, but is sensitive to the initial parameters and can be easily trapped in a local solution [3]. To develop the "route" in our optimization method, we first specify important data features (Focused Feature and Stabilized Feature), and define the optimization objective. In this work, the *optimization objective* was to increase the accuracy in fitting high frequency DBS data, while maintaining a reasonable accuracy in fitting low frequency DBS data. To realize this optimization objective, we designed a set of objective functions (Pushing Function and Stabilizing Function, see **Fig. 3B**) to emphasize high frequency DBS data, and balance the fitting accuracy among data across different DBS frequencies (5~200 Hz). By sequentially executing these objective functions, the route optimization method automatically navigates the exploiting Nelder-Mead optimizer to explore a route leading to the desired globally optimal solution that satisfies the optimization objective.

***Concatenate data from different frequencies of DBS***

We concatenated the experimental instantaneous firing rate of the data from different frequencies of Vim-DBS ({5, 10, 20, 30, 50, 100, and 200 Hz}). Such "concatenated experimental instantaneous firing rate" was the reference for the model fit. As well, we concatenated the rate network model simulations during the above frequencies of DBS, and computed the mean squared error (MSE) between the concatenated signals (model simulation and reference). The purpose of minimizing MSE based on concatenated signals was to fit the model consistently across different DBS frequencies.

To be more specific, we used the sampling resolution at 0.1 ms for all model simulations, and a simulated signal is denoted as $\boldsymbol{r}_{fq}(\Phi, \boldsymbol{t})$, which corresponds to a certain DBS frequency ($fq$) and parameter set $\Phi$ (Equation (1)); $\boldsymbol{r}_{fq}(\Phi, \boldsymbol{t}) = [\, r_{fq}(\Phi, t_1), \dots, r_{fq}\left(\Phi, t_{N_{fq}}\right)]$ and $N_{fq}$ is the total number of sampling time points. Similarly, the reference firing rate was sampled with the same resolution (0.1 ms), and was denoted as $\boldsymbol{P}_{fq}(\boldsymbol{t}) = [\, P_{fq}(t_1), \dots, P_{fq}\left(t_{N_{fq}}\right)]$; the corresponding MSE was computed as:

$$MSE_{fq}(\Phi) = \frac{1}{N_{fq}} \parallel \boldsymbol{r}_{fq}(\Phi, \boldsymbol{t}) - \boldsymbol{P}_{fq}(\boldsymbol{t}) \parallel^2 = \frac{1}{N_{fq}} \sum_{i=1}^{N_{fq}} [r_{fq}\,(\Phi\,, t_i) - P_{fq}(\,t_i)]^2 \qquad (16)$$

Finally, the MSE between concatenated signals was the weighted sum across different DBS frequencies:

$$MSE_{conc}(\boldsymbol{g}, \Phi) = \sum_{fq} g_{fq} * \ MSE_{fq}(\Phi) \qquad (17)$$

where $fq \in$ {5, 10, 20, 30, 50, 100, and 200 Hz}, $g_{fq}$ is the weight of $MSE_{fq}(\Phi)$, and $\boldsymbol{g} = [g_{5Hz}, \dots, g_{200Hz}]$ is the weight vector, which depends on the specific purposes in the process of our route optimization method.

***Data features and optimization objective functions***

Our *optimization goal* was to increase the accuracy in fitting data from high DBS frequencies (≥100 Hz), while maintaining a reasonable accuracy in fitting data from low DBS frequencies (<100 Hz). Thus, we stressed the data from high frequency of Vim-DBS, which is more clinically effective in treating essential tremor [8,9,10]. Based on our optimization goal, we defined two features of the Vim-DBS experimental instantaneous firing rate: Focused Feature and Stabilized Feature, whose differences were characterized by the weight vector $\boldsymbol{g}$ in computing MSE (Equation (17)). Focused Feature emphasized the high frequencies (100-Hz and 200-Hz) DBS data, and the corresponding weights ($g_{100Hz}$ and $g_{200Hz}$) were higher in computing the MSE (Equation (17), **Supplementary Table S3**). Stabilized Feature, namely, balanced the weights of high frequency and low frequency DBS data in the MSE computations; i.e., $g_{fq\leq 50Hz}$ was not far from $g_{fq\geq 100\mathrm{Hz}}$ (Equation (17), **Supplementary Table S3**). Focused Feature is important in increasing the fitting accuracy (MSE) to data from high frequency of DBS, and Stabilized Feature is critical in fitting consistent model parameters across all DBS frequencies.

In order to fit Focused Feature or Stabilized Feature, we designed two types of optimization objective functions: Pushing Function and Stabilizing Function, which were the MSE computations with different weight vectors $\boldsymbol{g}$ (Equation (17)). Pushing Function emphasizes the Focused Feature, whereas Stabilized Feature is stressed in Stabilizing Function. The output parameters from each optimization objective function were obtained with the MATLAB custom function "fminsearch", which implemented the Nelder-Mead simplex method [6,7]. Pushing Function (on Focused Feature) "pushes" the model parameters away from the local solution towards the parameter range better representing Focused Feature, then Stabilizing Function (on Stabilized Feature) "stabilizes" the model parameters by fitting consistently across the whole data (i.e., Stabilized Feature) with high accuracy. Such sequential execution of Pushing Function and Stabilizing Function (**Fig. 3B**) is the critical idea of the route optimization method in effectively finding the globally optimal solution. Essentially, the pushing-and-stabilizing strategy keeps updating and improving the prior knowledge of the initial model parameters for each iteration of fits.

***Preliminary fit***

The undetermined parameter set to be optimized was $\Phi = \{w_{ee}, w_{ie}, w_{ei}, w_{ii}, \tau_i, \tau_e, r_{E,b}, c, s, k\}$ (Equation (1)). The route optimization process started with a set of initial parameters $\Phi_{ini} = \{w_{ee,0}, w_{ie,0}, w_{ei,0}, w_{ii,0}, \tau_{i,0}, \tau_{e,0}, r_{E,b,0}, c_0, s_0, k_0\}$ (**Supplementary Table S4**), which were determined as follows: (i) The initial connectivity strength parameters ($\{w_{ee,0}, w_{ie,0}, w_{ei,0}, w_{ii,0}\}$) were all set to be 1; (ii) $\{\tau_{i,0}, \tau_{e,0}, r_{E,b,0}\}$ were from the anatomical and experimental literature [11,1,12,13,14,15]; (iii) $\{c_0, s_0, k_0\}$ were from fitting the reference firing rate with our previous single-ensemble model of the Vim neural group receiving DBS [2]. Starting from the initial parameter set $\Phi_{ini}$ , we did a preliminary model fit based on Stabilized Feature (i.e., stressing data from both low and high frequencies of DBS), and the weight of each DBS frequency was given by the vector $\boldsymbol{g}_{pre}$ (Equation (17), **Supplementary Table S3**). The output parameter set of this preliminary fit was denoted as $\Phi_{pre}$ (**Supplementary Table S4**). The total fitting error (ER, Equation (2)) with $\Phi_{ini}$ and $\Phi_{pre}$ were very large (>40%, **Supplementary Table S5**), and model simulations with these two parameter sets were very different from the experimental data.

***Main stages of the route optimization***

The preliminary fit result $\Phi_{pre}$ entered the main stages of the route optimization: Global Stage and Refining Stage (**Fig. 3**). Global Stage explores the parameter space and finds the appropriate parameter range with low total fitting error (ER, Equation (2)); Refining Stage exploits the "low error area" attained by Global Stage, and finds the optimal parameter set $\Phi_{optimal}$ that minimizes the total fitting error (**Fig. 3A**). Compared with Global Stage, Refining Stage stresses Focused Feature to a higher extent (**Supplementary Table S3**). This design was compatible with our optimization goal in increasing the fitting accuracy of data from high frequencies of Vim-DBS (more clinically effective). Global Stage and Refining Stage comprise iterations of sequentially executed optimization objective functions – Pushing Function and Stabilizing Function (**Fig. 3B**). Pushing Function in Global Stage (respectively, Refining Stage) was denoted as $J_{G,P}(\Phi)$ (respectively, $J_{R,P}(\Phi)$), and Stabilizing Function in Global Stage (respectively, Refining Stage) was denoted as $J_{G,S}(\Phi)$ (respectively, $J_{R,S}(\Phi)$); these four optimization objective functions were defined as follows:

$$J_{G,P}(\Phi) = MSE_{conc}(\boldsymbol{g}_{G,P}, \Phi) = \sum_{fq} g_{G,P,fq} * \ MSE_{fq}(\Phi) \qquad (18)$$

$$J_{R,P}(\Phi) = MSE_{conc}(\boldsymbol{g}_{R,P}, \Phi) = \sum_{fq} g_{R,P,fq} * MSE_{fq}(\Phi) \tag{19}$$

$$J_{G,S}(\Phi) = MSE_{conc}(\boldsymbol{g}_{G,S}, \Phi) = \sum_{fq} g_{G,S,fq} * MSE_{fq}(\Phi) \tag{20}$$

$$J_{R,S}(\Phi) = MSE_{conc}(\boldsymbol{g}_{R,S}, \Phi) = \sum_{fq} g_{R,S,fq} * MSE_{fq}(\Phi) \tag{21}$$

where $\Phi$ is the undetermined parameter set (Equation (1)); $\boldsymbol{g}_{G,P}$ , $\boldsymbol{g}_{R,P}$ , $\boldsymbol{g}_{G,S}$ and $\boldsymbol{g}_{R,S}$ are the corresponding weight vector (**Supplementary Table S3**); $fq$ is the DBS frequency and $fq \in$ {5, 10, 20, 30, 50, 100, and 200 Hz}; MSE was computed by Equations (9) and (10).

In the 1st iteration of Global Stage, starting from $\Phi_{pre}$ (the preliminary fit result), we obtained $\Phi_{F,W_{pre}}$ by optimizing $J_{G,P}(\Phi)$ (Equation (18)); in $\Phi_{F,W_{pre}}$ the subscript "F" represents that it is the output of a Pushing Function $J_{G,P}(\Phi)$ emphasizing Focused Feature, and the subscript "$W_{pre}$" stresses the connectivity strength features (Equation (1)). Then, starting from $\Phi_{F,W_{pre}}$ , we optimized $J_{G,S}(\Phi)$ (Equation (20)) and obtained $\Phi_{S,W_1}$ . In $\Phi_{S,W_1}$ , the subscript "S" represents that it is the output of a Stabilizing Function $J_{G,S}(\Phi)$ emphasizing Stabilized Feature, and the subscript "$W_1$" shows that $\Phi_{S,W_1}$ is the output of 1st iteration of Global Stage. To conclude, the 1st iteration of Global Stage is summarized as $\Phi_{pre} \xrightarrow{J_{G,P}(\Phi)} \Phi_{F,W_{pre}} \xrightarrow{J_{G,S}(\Phi)} \Phi_{S,W_1}$. The $k^{\text{th}}$ iteration $(k \geq 2)$ of Global Stage is summarized as $\Phi_{S,W_{k-1}} \xrightarrow{J_{G,P}(\Phi)} \Phi_{F,W_{k-1}} \xrightarrow{J_{G,S}(\Phi)} \Phi_{S,W_k}$ . Note that when optimizing with Pushing Function $J_{G,P}(\Phi)$, the connectivity strength matrix $W_{k-1}$ was always fixed. The purpose of this design was to increase the prediction robustness of the connectivity strength, which was the essential feature of a neural network.

The end of Global Stage is denoted as $\Phi_{S,W_G}$ , which is also the start of Refining Stage. The $k^{\text{th}}$ iteration $(k \geq 1)$ of Refining Stage is summarized as $\Phi_{S,W_{G+k-1}} \xrightarrow{J_{R,P}(\Phi)} \Phi_{F,W_{G+k-1}} \xrightarrow{J_{R,S}(\Phi)} \Phi_{S,W_{G+k}}$ , where $J_{R,P}(\Phi)$ (Equation (19)) is the Pushing Function, and $J_{R,S}(\Phi)$ is the Stabilizing Function (Equation (21)). The final output of Refining Stage was the globally optimized model parameter set $\Phi_{optimal}$ with minimized fitting errors, which satisfied the optimization *exit rule*:

$$\text{ER} \leq 8\%, NMSE\ (100\ Hz) + NMSE\ (200\ Hz) \leq 9\% \tag{22}$$

where ER is the "total fitting error" (Equation (2)), and NMSE represents "normalized mean squared error". See **Supplementary Tables S4 and S5** for the evolution of the model parameter set $\Phi$ and its fitting errors during the route optimization process, from the initial parameter set ($\Phi_{ini}$) to the globally optimal solution ($\Phi_{optimal}$).

***Generalizability to other datasets***

Our route optimization method can be implemented in a general setting, comprising a biological dataset and an appropriate network model. We first identify the physiological features of the dataset to be studied, and specify the optimization objective. In many scenarios, we can identify the dataset as consisting of two physiological features: one prominent feature that we currently focus on, and another secondary feature that may also be important. When fitting a network model to such a dataset, we can define its prominent feature as Focused Feature, define the weight-balance between its two features as Stabilized Feature, and proceed with the "pushing and stabilizing" strategy in this work (**Fig. 3**) to design a route leading to the globally optimal model parameters that minimize fitting errors. There are also other scenarios wherein a different composition of physiological features – e.g., two prominent features or three features – is appropriate for a specific dataset. In such scenarios, we can design the optimization route accordingly. For example, when fitting two prominent features, the definition of Focused Feature can alternate between the two features during the route optimization process. When fitting multiple (≥3) features, we can stress each Focused Feature sequentially, while maintaining robustness by defining the weight-balance among all features as Stabilized Feature. Thus, the design of the route optimization is highly flexible to specific datasets and implementation goals. Compared with the traditional optimization methods – parameter hand tuning or one-step optimization (e.g., *fminsearch* in Matlab [6,7]), we anticipate that the route optimization method can be highly effective in finding the globally optimal solution – which characterizes physiological underpinnings – in a relatively high dimensional model parameter space.

#### Supplementary Notes 2– further discussions

##### *DBS impact as an external drive*

DBS impacts the firing rate dynamics as an external drive of the Vim-network (**Fig. 6**). In low-frequency Vim-DBS (5∼50 Hz), we observed large evoked activities in response to each DBS pulse (**Fig. 2**). In order to clarify the underlying mechanism, we tracked the evolution of mean firing rate in inter-DBS-pulse-intervals for each recorded frequency of DBS (5∼200 Hz) (**Fig. 6A – D**); the mean firing rate is directly related to the metabolic energy consumed in physiological processes [57]. In **Fig. 6A – D**, the firing rate at DBS pulse $\#i$ ($i \geq 1$) is defined as the mean firing rate in the $[(i-1)^{th}, (i+1)^{th}]$ inter-pulse-intervals, where "$0^{th}$" represents the baseline firing rate before the onset of DBS; this baseline firing rate is used to define the firing rate at DBS pulse #0. During the DBS pulses of each frequency, we showed the firing rate dynamics of the Vim neurons in the experimental data (**Fig. 6A**), the Hebbian network (**Fig. 6B**), and the BA network (**Fig. 6C**). In **Fig. 6D**, we showed the firing rate dynamics of the external inhibitory nuclei in the BA network. We observed that the firing rate is closed to the steady-state response after receiving energy delivered by ∼60 DBS pulses, across different DBS frequencies. We also observed that the BA network accurately tracked the firing rate of Vim-DBS during each DBS frequency (5 to 200 Hz) (**Fig. 6A and C**). However, the Hebbian network deviated from the experimental data, specifically for high-frequency DBS data (100 and 200 Hz). Both amplitude and duration of the initial large transient response (the initial ∼60 DBS pulses) were not accurately estimated (**Fig. 6A and B**).

##### *Developing macroscopic network models and control systems*

We developed a rate network model and implemented it in the neural circuits impacting thalamic Vim neurons receiving DBS. To implement the model in improving the clinical effects of DBS, it is essentially important to develop a closed-loop DBS control system that automatically adjusts the stimulation parameters to best suit each individual patient [71,72]. The closed-loop DBS control system consists of a biomarker as the feedback signal (e.g., oscillations in local field potentials (LFP) [71], power of electromyography (EMG) [72]), and a computational model that predicts the clinical state from such recordings in order to control

DBS in a feedback-driven manner [73]. For example, in treating Parkinson's disease or essential tremor with closed-loop DBS control, the computational model can be a macroscopic network model of the disease-related physiological mechanism underlying the thalamocortical and cerebellar circuits [73,74]. In this work, we investigated the Vim-network with our model, and accurately reproduced the experimental recordings in Vim neurons across various DBS frequencies (5 to 200 Hz). For further work, we can propagate the DBS impact on Vim neurons out towards their efferent target circuits, and build a macroscopic thalamocortical-cerebellar network model that quantifies the neurological disease mechanism. Finally, we will construct a closed-loop DBS control system based on the macroscopic thalamocortical-cerebellar network model, and clinically treat patients with neurological disorders by automatically adjusting DBS frequencies in continuous time.

***Relationship with other network models***

Dynamics in neural networks emerge from the complex interplay among a large number of neurons. The spiking network model is a direct simulation of the interaction among neurons, and the advantage of such a model lies in detecting neural activity patterns based on microscopic-level dynamics, including the membrane potential, activation probability of ion channels, and strength and variability of inter-neuron connections [22,77,40]. Typical examples of the spiking network model include leaky integrate-and-fire (LIF) [78], Izhikevich [79] and Hodgkin & Huxley types [77]. Despite the success of the spiking network model, its complexity and high dimensionality preclude the theoretical interpretation of network-level mechanisms [40,26].

The mean-field firing rate network model of neural populations has been the mainstay for interpreting mechanisms on the neural circuit level [40], including the E/I balance [24], connectivity among brain regions [80] and large-scale oscillations [74]. The typical and classical mean-field firing rate network model is of the Wilson-Cowan type [81], which assumes the homogeneity of neurons in one population [26,40]. Compared with the spiking network model, the mean-field firing rate network model reduces the complexity and dimensionality, and is more robust when tracking population-level dynamics and interactions [26,40]. Besides, the mean-field firing rate network model has fewer parameters, and is thus more efficient for implementation [73] and parameter automatic optimization [82].

Typically, a mean-field model assumes infinitely large populations, and is less sensitive to abrupt changes in inputs to local neurons [40]. In this work, we developed a mean-field firing rate network model that is consistent with the first principles, and also responds effectively to abrupt local input changes, i.e., the injection of DBS pulses to a local group of Vim neurons. We modeled the immediate impact of DBS pulses to the local Vim neural group as inducing synaptic release, and the synapses were characterized with the Tsodyks & Markram (TM) model [36]; such modeling of DBS effects is consistent with previously established works [12,13,83]. Our whole model consists of the local neural group directly receiving DBS, and its interactions with the external excitatory and inhibitory neural groups; the three neural groups are mutually recurrent (**Fig. 1**).

***Principle of identifying neural networks***

With our rate network model, we explored the physiological mechanism underlying Vim-network in response to DBS. We found that the Vim-network can be identified as an inhibition-stabilized network with a Balanced Amplification mechanism, in which the strong external input and recurrent excitation are balanced by equally strong feedback inhibition. The Balanced Amplification mechanism (represented by the optimal model parameters) accurately fitted the experimental data, and was consistent across various DBS frequencies (5~200 Hz) (**Fig. 2**, **Supplementary Table S7**). In contrast, the Hebbian mechanism (weak inhibition) was only appropriate for DBS data with low stimulation frequency (≤50 Hz), and missed the features of neural responses to high-frequency (≥100 Hz) DBS (**Supplementary Fig. S4**, **Supplementary Table S7**). Thus, in order to more fully identify the physiological components of the Vim-network, we need to fit the model consistently across data from a wide spectrum of DBS frequencies; if we confine the model fit within a small range of DBS frequencies, we probably identify the network mechanism erroneously. The same principle can be generalized to different types of neuromodulations: To identify the mechanism of a neural network with neuromodulation, we need to observe and model the neural response to a full range of the neuromodulation parameter, e.g., the frequency, pulse duration, or intensity of injected currents [75], focused ultrasound [76] and visual stimuli [34].

### Supplementary figures

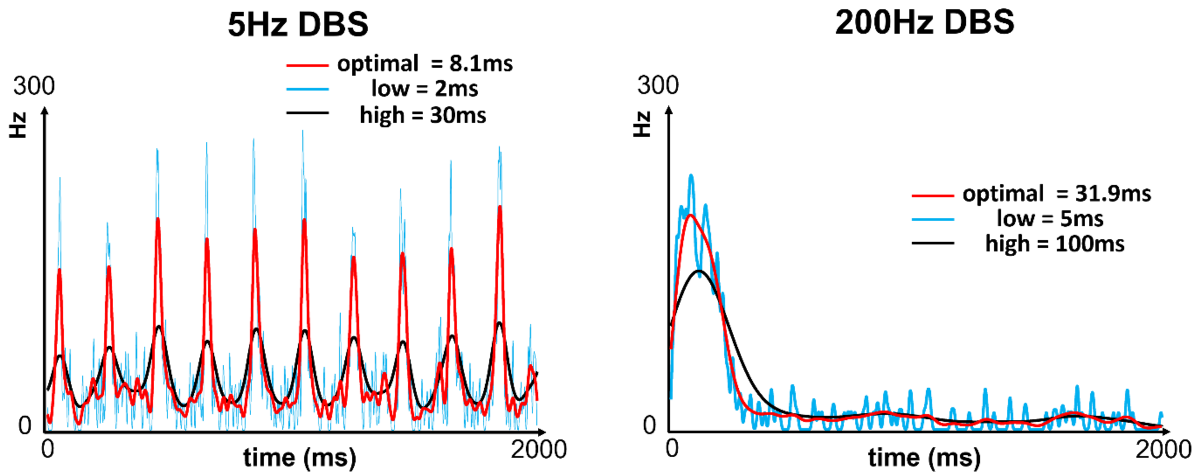

**Supplementary Fig. S1. Compare different Gaussian time histogram kernel widths for computing instantaneous firing rate from DBS experimental data**

*We compute the instantaneous firing rate using a time histogram method from the experimental data recorded in the ventral intermediate nucleus (Vim) during Vim-DBS with various stimulation frequencies; we take 5-Hz and 200-Hz DBS data as the examples. The instantaneous firing rate is computed by time histogram with a Gaussian kernel, and the optimal kernel width is obtained by the method in Shimazaki and Shinomoto (2007) and (2010)* [16,17]*. We compare the instantaneous firing rate from the optimal kernel with the result from a low and a high kernel, respectively.*

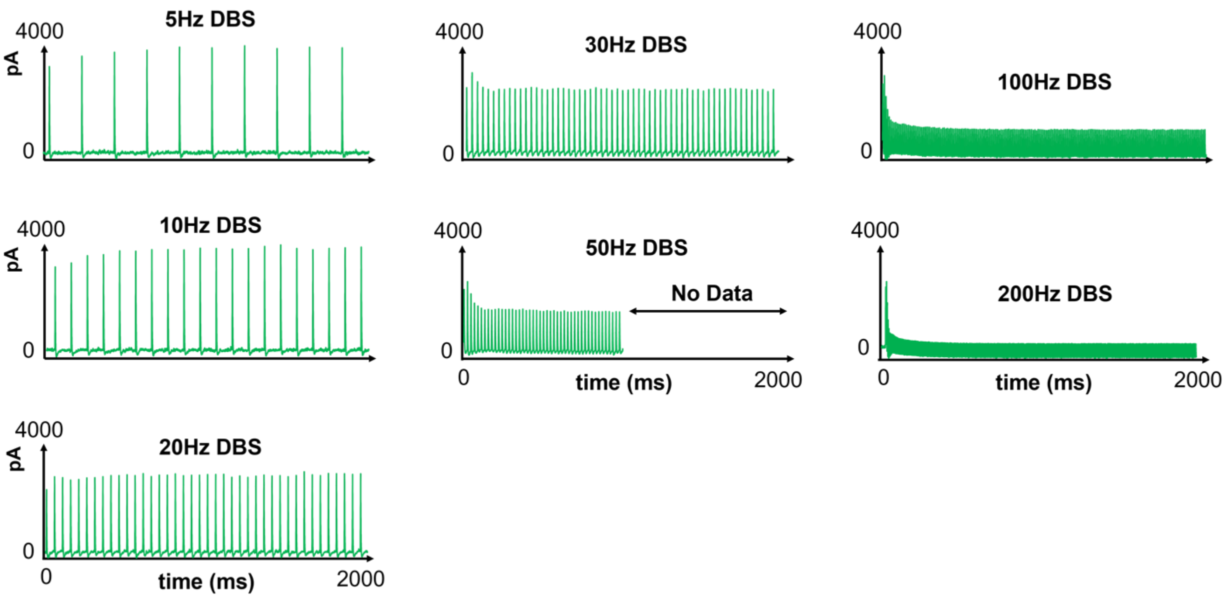

**Supplementary Fig. S2. Modeled DBS-induced postsynaptic current into Vim ($I_{DBS}$)**

*We model the DBS-induced postsynaptic current ($I_{DBS}$, Equation (1)) into a neuron in the ventral intermediate nucleus (Vim), with the Tsodyks & Markram model [18] of short-term synaptic plasticity (STP) (**Methods S3**).We present the modeled current during Vim-DBS of various stimulation frequencies (5~200 Hz).*

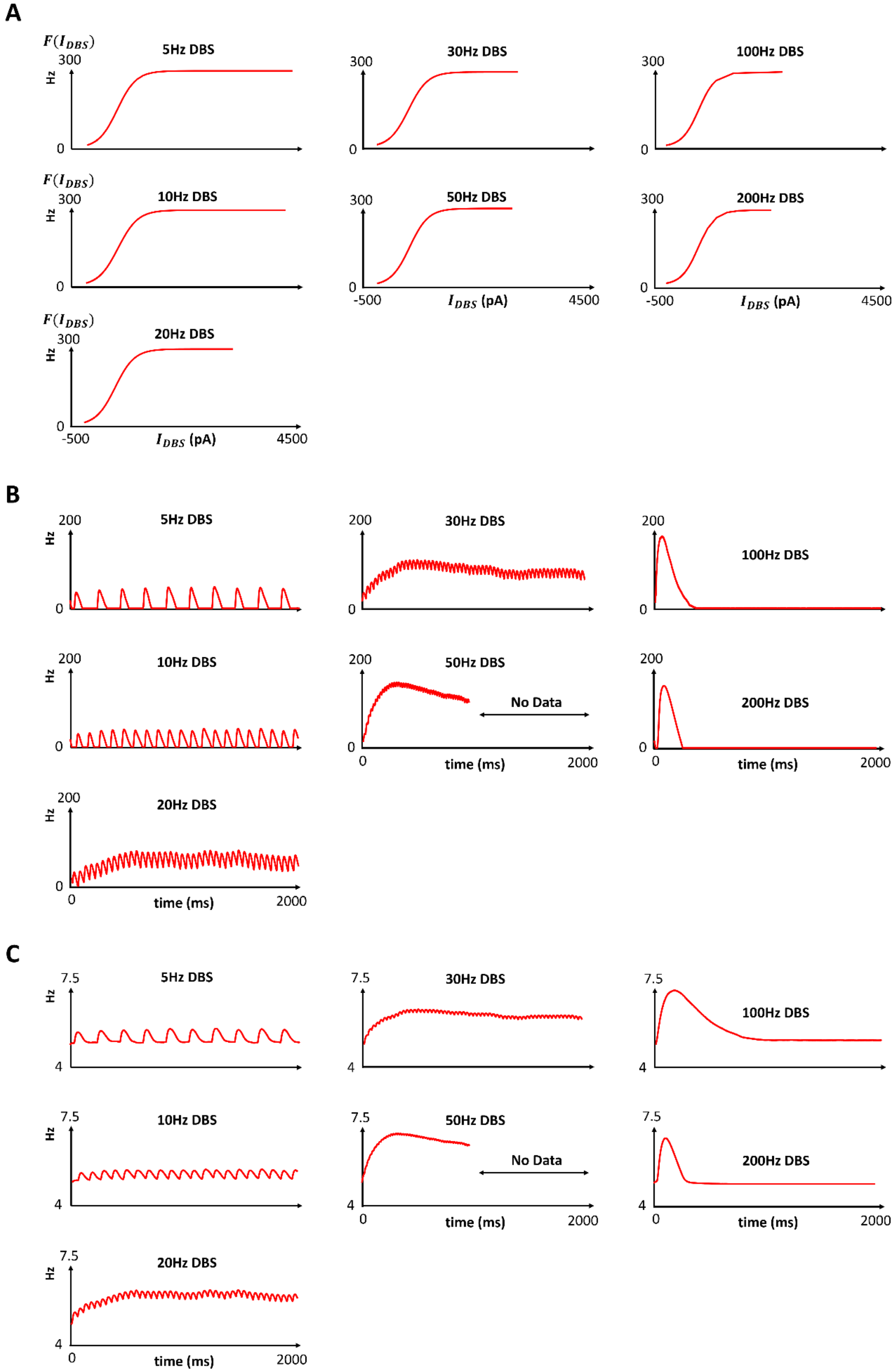
A
5Hz DBS
30Hz DBS
100Hz DBS
10Hz DBS
50Hz DBS
200Hz DBS
20Hz DBS
$F(I_{DBS})$
Hz
300
0
-500
4500
$I_{DBS}$ (pA)
B
200
No Data
time (ms)
2000
C
7.5
4

#### Supplementary Fig. S3. Optimal rate network model fit result of Vim-DBS human data ($\boldsymbol{F(I_{DBS})}$, $\boldsymbol{r_E}$ and $\boldsymbol{r_I}$)

*We present simulated variables ($F(I_{DBS})$, $r_E$ and $r_I$) corresponding to the optimal rate network model fit (**Fig. 2**), during DBS in the ventral intermediate nucleus (Vim), with various stimulation frequencies (5~200 Hz). See Equation (1) for the details of these variables. The optimal rate network model fit is characterized by the Balanced Amplification (BA) mechanism (**Fig. 4C**).*

*(A). Firing rate non-linear variability corresponding to $I_{DBS}$ (see **Supplementary Fig. S2**) through a sigmoid transfer function ($F(I_{DBS})$).*

*(B) Firing rate of the external excitatory nuclei ($r_E$).*

*(C) Firing rate of the external inhibitory nuclei ($r_I$).*

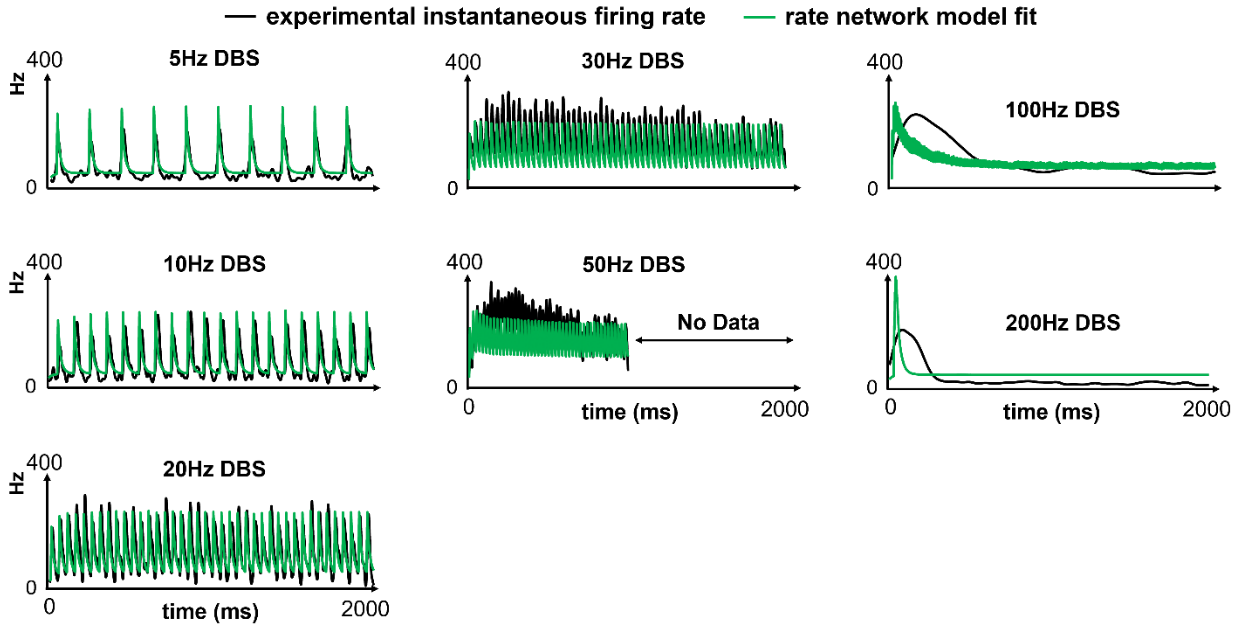

**Supplementary Fig. S4. Hebbian rate network model fit result of Vim-DBS human data ($r_D$)**

*We present the rate network model fit result with the Hebbian mechanism (**Fig. 4C** and **Supplementary Table S4**), during DBS in the ventral intermediate nucleus (Vim), with various stimulation frequencies (5~200 Hz).*

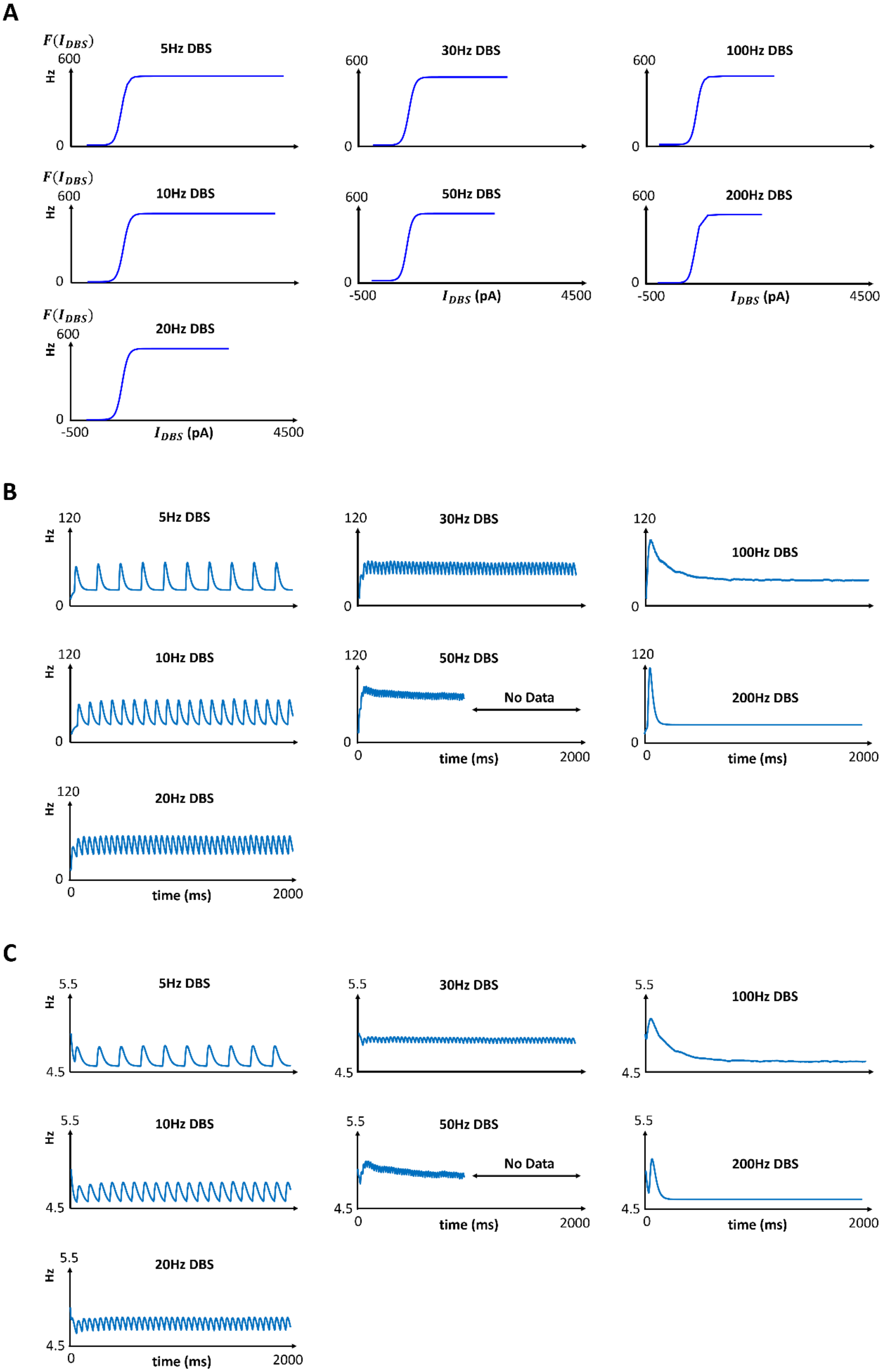

A
$F(I_{DBS})$
600
Hz
0
5Hz DBS
10Hz DBS
20Hz DBS
30Hz DBS
50Hz DBS
100Hz DBS
200Hz DBS
-500
$I_{DBS}$ (pA)
4500
B
120
time (ms)
2000
No Data
C
5.5
4.5

#### Supplementary Fig. S5. Hebbian rate network model fit result of Vim-DBS human data ($\boldsymbol{F(I_{DBS})}$, $\boldsymbol{r_E}$ and $\boldsymbol{r_I}$)

*We present simulated variables ($F(I_{DBS})$, $r_E$ and $r_I$) corresponding to the rate network model with the Hebbian mechanism (**Supplementary Fig. S4**, **Fig. 4C** and **Supplementary Table S4**), during DBS in the ventral intermediate nucleus (Vim), with various stimulation frequencies (5~200 Hz). See Equation (1) for the details of these variables.*

*(A). Firing rate non-linear variability corresponding to $I_{DBS}$ (see **Supplementary Fig. S2**) through a sigmoid transfer function ($F(I_{DBS})$).*

*(B) Firing rate of the external excitatory nuclei ($r_E$).*

*(C) Firing rate of the external inhibitory nuclei ($r_I$).*

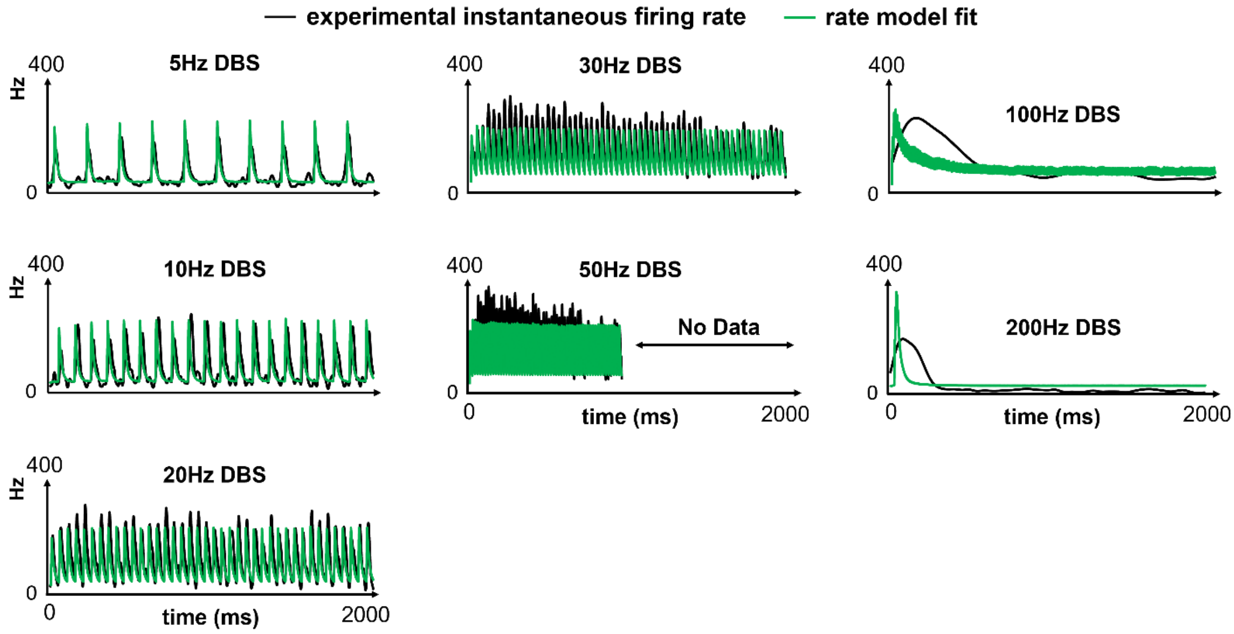

**Supplementary Fig. S6. Single-ensemble rate model fit result of Vim-DBS human data**

*We present the single-ensemble rate model [2] fit result, during DBS in the ventral intermediate nucleus (Vim), with various stimulation frequencies (5~200 Hz). This rate model was solely of the Vim neural group directly receiving DBS, and ignored recurrent connections with other nuclei [2]. In this single-ensemble rate model fit, we restrict the time constant of the Vim neurons (excitatory) to be ≤15 ms, to be consistent with the physiological literature [1,19,20].*

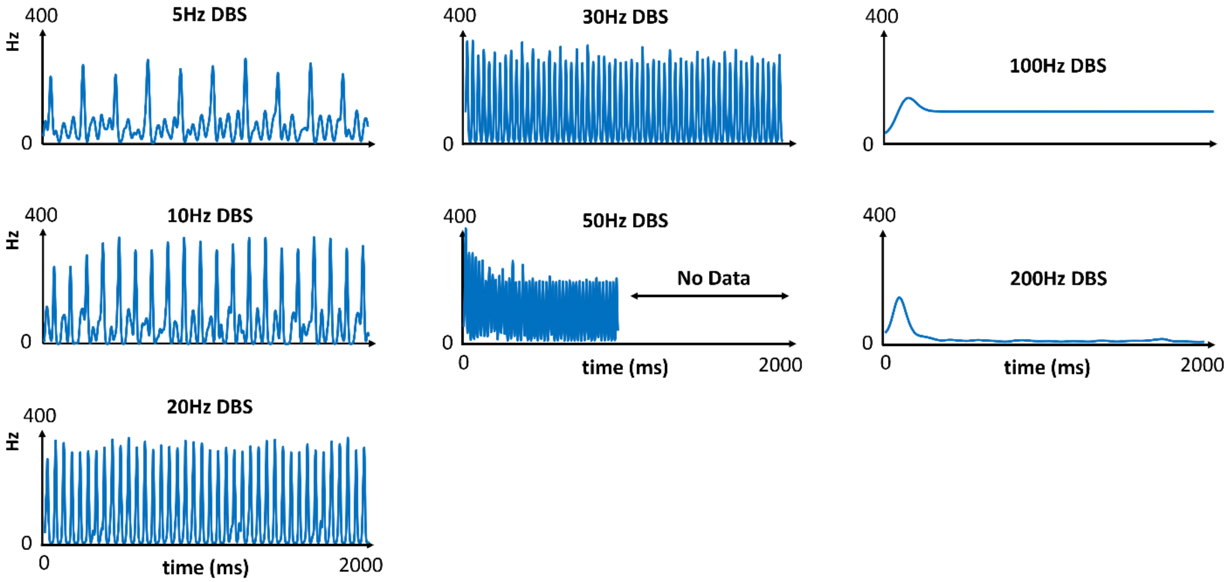

#### Supplementary Fig. S7. Izhikevich spiking network model simulated firing rate of Vim neurons during Vim-DBS (Hebbian mechanism)

*We simulate an Izhikevich spiking network model during DBS in the ventral intermediate nucleus (Vim), with various stimulation frequencies (5~200 Hz) (**Methods S6**). For each frequency of DBS, the instantaneous firing rate is computed by a time histogram with Gaussian kernel width shown in **Supplementary Table S1**. In this plot, the spiking network model is with the Hebbian mechanism, characterized by the corresponding connectivity strength matrix W (Equation (1), **Supplementary Table S4**).*

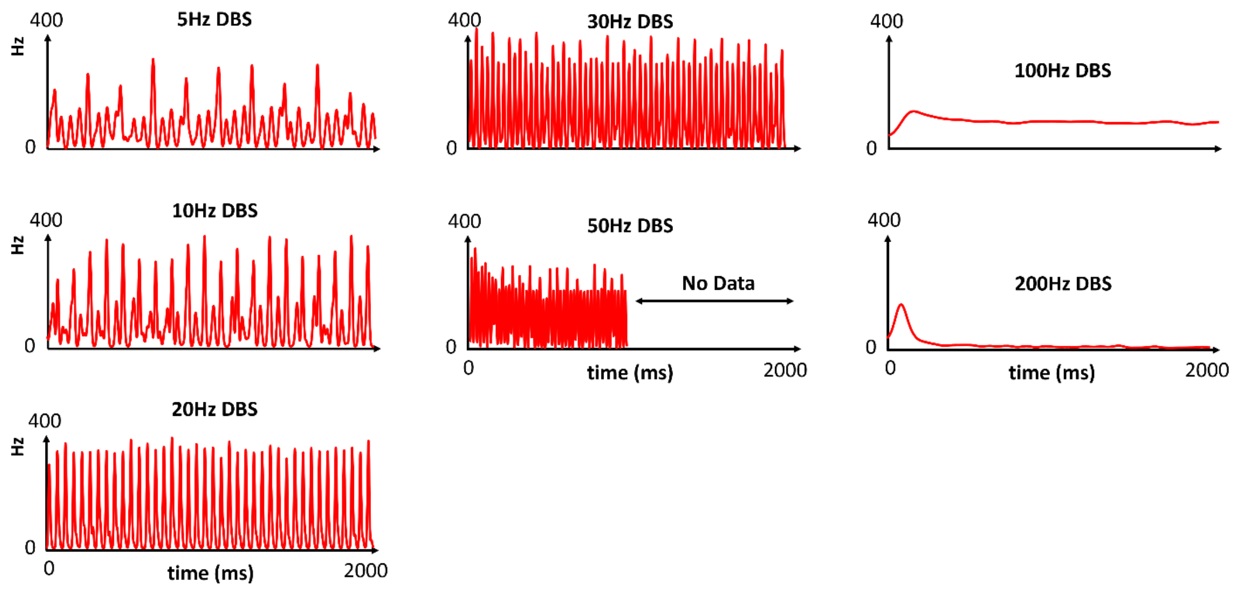

#### Supplementary Fig. S8. Izhikevich spiking network model simulated firing rate of Vim neurons during Vim-DBS (Balanced Amplification mechanism)

*We simulate an Izhikevich spiking network model during DBS in the ventral intermediate nucleus (Vim), with various stimulation frequencies (5~200 Hz) (**Methods S6**). For each frequency of DBS, the instantaneous firing rate is computed by a time histogram with Gaussian kernel width shown in **Supplementary Table S1**. In this plot, the spiking network model is with the Balanced Amplification (BA) mechanism, characterized by the corresponding connectivity strength matrix W (Equation (1), table in **Fig. 2**).*

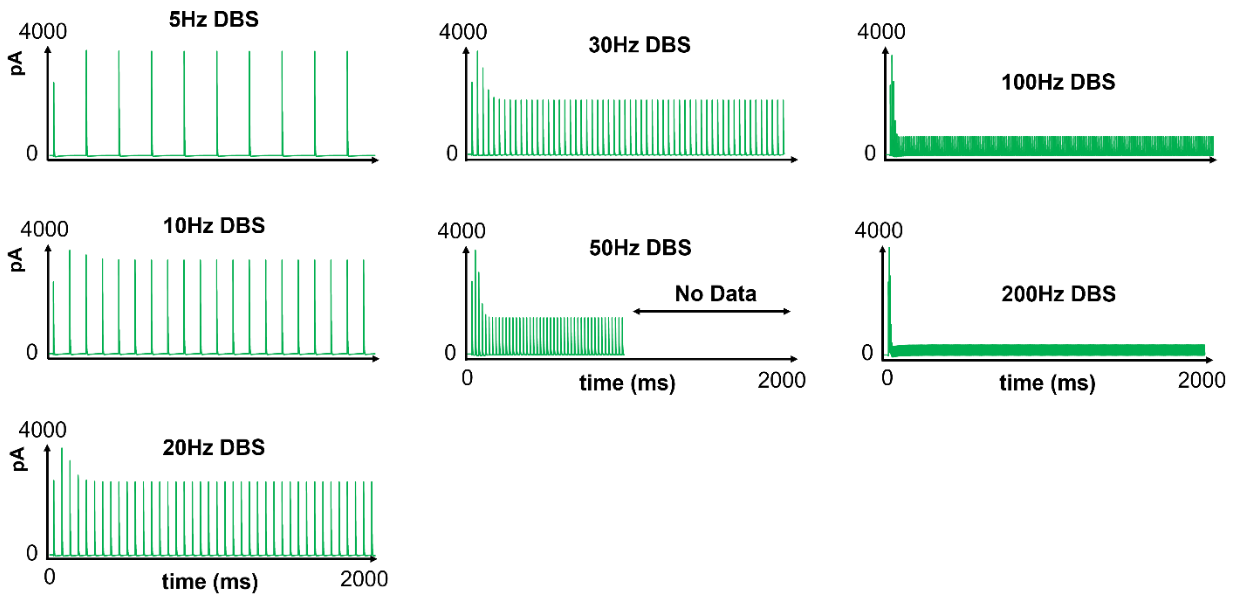

#### Supplementary Fig. S9. Tsodyks & Markram model used in the Izhikevich spiking network model

*We present the simulation results of the Tsodyks & Markram model [21,22] used in "tsodyks2_synapse", which is a built-in function of the Neural Simulation Tool (NEST) platform [23]. "tsodyks2_synapse" is used in the Izhikevich spiking network model (**Methods S6**), and its parameters are specified in **Supplementary Table S9**. We present the simulations of "tsodyks2_synapse" in response to different DBS frequencies (5~200 Hz). The simulation of "tsodyks2_synapse" used in the spiking model is consistent with the Tsodyks & Markram model used in the rate model (**Supplementary Fig. S2**).*

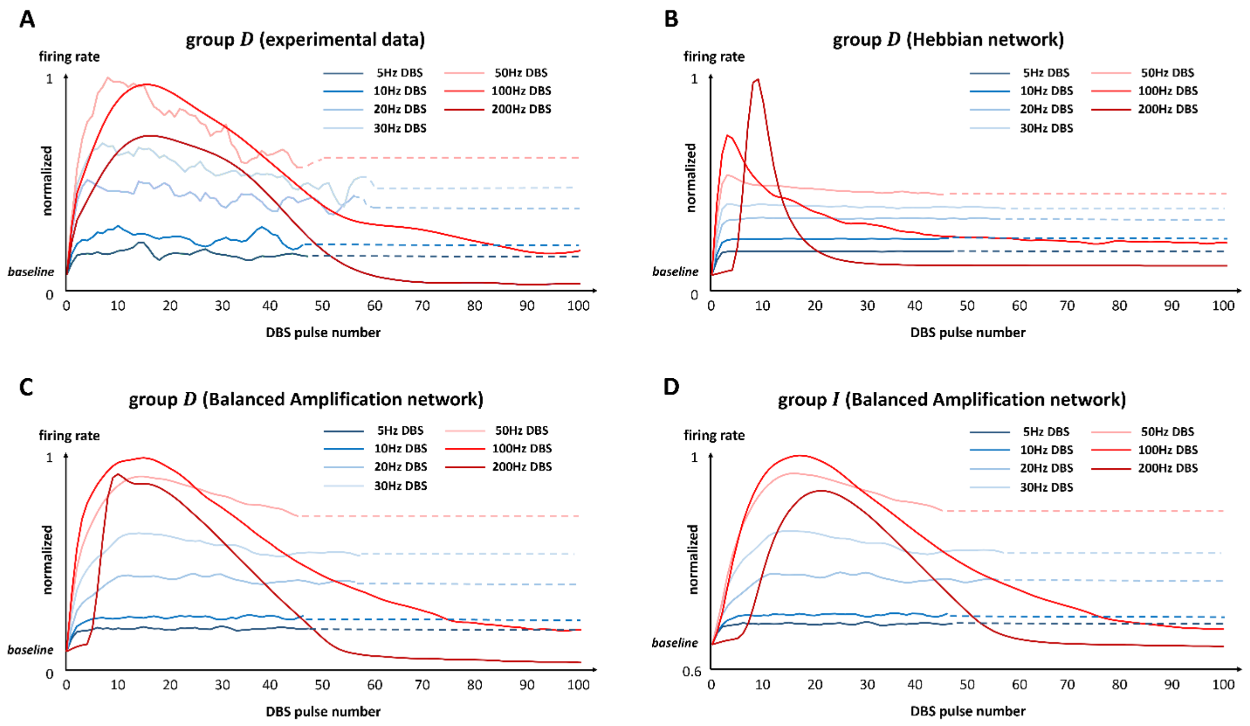

#### Supplementary Fig. S10. DBS impact as an external drive during different stimulation frequencies

*In each plot, the firing rate at DBS pulse #i* ($i \geq 1$) *is defined as the mean firing rate in the* $[(i-1)^{th}, (i+1)^{th}]$ *inter-pulse-intervals, where "*$0^{th}$*" represents the baseline firing rate before the onset of DBS; this baseline firing rate is used to define the firing rate at DBS pulse #0. The solid lines represent data and fits during clinical recordings, and the dashed lines represent the steady state estimations after clinical recordings. Note that in low-frequency (≤50 Hz) DBS data and fits during clinical recording, the maximal number of DBS pulses is smaller, because the inter-pulse-interval is longer, and the total length of recording is restricted (≤10 s for each recording). In each plot, the firing rate is normalized to the maximum rate across all DBS frequencies.*

*(A) Firing rate dynamics of the experimental recordings in the Vim neurons directly receiving DBS (group D, see* ***Fig. 1****).*

*(B) Firing rate dynamics of the model simulations with the Hebbian mechanism, simulated on the Vim neurons directly receiving DBS.*

*(C) Firing rate dynamics of the model simulations with the Balanced Amplification mechanism and the optimal model parameters (see* ***Fig. 2****), simulated on the Vim neurons directly receiving DBS.*

*(D) Firing rate dynamics of the model simulations with the Balanced Amplification mechanism and the optimal model parameters (see* ***Fig. 2****), simulated on the inhibitory nuclei (group I, see* ***Fig. 1****).*

### Supplementary tables

| DBS frequency | 5Hz | 10Hz | 20Hz | 30Hz | 50Hz | 100Hz | 200Hz |
|---|---|---|---|---|---|---|---|
| Gaussian kernel(ms) | 8.1 | 5.7 | 4.4 | 3.9 | 3.5 | 53.4 | 31.9 |

##### Supplementary Table S1. The optimized Gaussian time histogram kernel for spiking data from each DBS frequency

*We show the optimized Gaussian time histogram kernel (i.e., standard deviation) for computing the instantaneous firing rate of the experimental spike trains recorded during each frequency of DBS. The optimized Gaussian kernel is obtained by the methods in Shimazaki and Shinomoto (2007) and (2010)* [16,17].

| ***excitatory synapses*** **$N_{exc}$ = 450; $w_{exc}$ = 54.25** | | | | | | ***inhibitory synapses*** **$N_{inh}$ = 50; $w_{inh}$ = 93.00** | | | | | |
|---|---|---|---|---|---|---|---|---|---|---|---|
| **para** / **type** | $U$ | $\tau_{facil}$ (ms) | $\tau_{rec}$ (ms) | $\tau_s$ (ms) | $A$ | **para** / **type** | $U$ | $\tau_{facil}$ (ms) | $\tau_{rec}$ (ms) | $\tau_s$ (ms) | $A$ |
| **F (40%)** | 0.19 | 670 | 138 | | | **F (40%)** | 0.016 | 376 | 45 | | |
| **P (20%)** | 0.45 | 326 | 329 | 2 | 1 | **P (30%)** | 0.29 | 62 | 144 | 8.5 | 1 |
| **S (40%)** | 0.04 | 17 | 85 | | | **S (30%)** | 0.25 | 21 | 706 | | |

##### Supplementary Table S2. Tsodyks & Markram model parameters of synapses projected to one Vim neuron

*We show the parameters related to the Tsodyks & Markram model* [18]*, which is implemented to compute the DBS-induced post-synaptic current ($I_{DBS}$ , Equation (5)) into the Vim neurons directly receiving DBS; "Vim" represents "thalamic ventral intermediate nucleus". For one Vim neuron, we model that it receives inputs from 500 synapses, with 90% excitatory synapses ($N_{exc}$ = 450) and 10% inhibitory synapses ($N_{inh}$ = 50). $w_{exc}$ and $w_{inh}$ are the scaling weights of the post-synaptic excitatory ($I_{exc}$) and inhibitory ($I_{inh}$) currents (Equation (5)). Both excitatory and inhibitory synapses consist of 3 types: facilitation ("F"), pseudo-linear ("P"), and depression ("S"). For excitatory synapses, "F (40%)" represents "the facilitation type of synapses accounted for 40% of all the excitatory synapses"; similar meanings for other synaptic types, and the inhibitory synapses. "para" represents "the Tsodyks & Markram model parameters", which consist of "U" (scaling factor), "$\tau_{facil}$" (facilitation time constant), "$\tau_{rec}$" (recovery time constant), "$\tau_s$" (post-synaptic time constant) and "A" (absolute response amplitude) (Equations (6) – (8)).*

| | 5Hz | 10Hz | 20Hz | 30Hz | 50Hz | 100Hz | 200Hz |
|---|---|---|---|---|---|---|---|
| $\boldsymbol{g}_{G,S}$ and $\boldsymbol{g}_{pre}$ | 24 | 0.6 | 1.2 | 1 | 3.6 | 7.2 | 12.75 |
| $\boldsymbol{g}_{R,S}$ | 24 | 0.6 | 1.2 | 1 | 3.6 | 7.2 | 45 |
| $\boldsymbol{g}_{G,P}$ | 6 | 3 | 7.5 | 1 | 6 | 24.6 | 157.5 |
| $\boldsymbol{g}_{R,P}$ | 6 | 3 | 7.5 | 1 | 6 | 24.6 | 375 |

#### Supplementary Table S3. Weight vectors of data from different frequencies of DBS

*We show the weight vectors* $\boldsymbol{g}$ *for the model fit MSE (mean squared error) to data from each DBS frequency;* $\boldsymbol{g}$ *is used to compute the concatenated MSE across different DBS frequencies (Equation (17)). All the weights are scaled by the weights of 30-Hz DBS data (normalized to 1). "*$\boldsymbol{g}_{pre}$*" represents the weight vector used in the preliminary model fit (**Supplementary Notes**) "*$\boldsymbol{g}_{G,S}$*" represents the weight vector used in Stabilizing Function (stressing Stabilized Feature) of Global Stage of the route optimization, and* $\boldsymbol{g}_{G,S} = \boldsymbol{g}_{pre}$ *; "*$\boldsymbol{g}_{R,S}$*" corresponds to Stabilizing Function of Refining Stage (**Supplementary Notes**). "*$\boldsymbol{g}_{G,P}$*" corresponds to Pushing Function (stressing Focused Feature) of Global Stage, and "*$\boldsymbol{g}_{R,P}$*" corresponds to Pushing Function of Refining Stage (**Supplementary Notes**).*

| | $w_{ee}$ | $w_{ie}$ | $w_{ei}$ | $w_{ii}$ | $\tau_i$ (ms) | $\tau_e$ (ms) | $r_{E,b}$ (Hz) | $c$ | $s$ | $k$ |
|---|---|---|---|---|---|---|---|---|---|---|
| $\Phi_{ini}$ | 1.00 | 1.00 | 1.00 | 1.00 | 13.0 | 5.00 | 40.0 | 324 | 7.51 $\times 10^{-3}$ | 360 |
| $\Phi_{pre}$ | 0.447 | 0.365 | 0.585 | 0.197 | 18.8 | 9.40 | 20.9 | 465 | 9.98 $\times 10^{-3}$ | 616 |
| $\Phi_{Hebbian}$ (= $\Phi_{(G,1)}$) | 0.282 | 3.01 $\times 10^{-3}$ | 0.764 | 0.147 | 24.0 | 9.40 | 10.1 | 562 | 1.45 $\times 10^{-2}$ | 584 |
| $\Phi_{(G,n_2)}$ | 0.390 | 1.00 $\times 10^{-3}$ | 2.41 | 1.02 $\times 10^{-3}$ | 17.9 | 8.65 | 10.0 | 418 | 1.77 $\times 10^{-2}$ | 577 |
| $\Phi_{(G,n_3)}$ | 0.437 | 1.28 $\times 10^{-3}$ | 3.59 | 1.07 $\times 10^{-3}$ | 18.8 | 9.40 | 12.5 | 371 | 3.68 $\times 10^{-2}$ | 571 |
| $\Phi_{(G,n_4)}$ | 0.492 | 1.73 $\times 10^{-3}$ | 6.31 | 1.00 $\times 10^{-3}$ | 17.4 | 5.05 | 10.0 | 227 | 5.18 $\times 10^{-3}$ | 469 |
| $\Phi_{(R,n_5)}$ | 0.487 | 1.00 $\times 10^{-3}$ | 8.67 | 1.00 $\times 10^{-3}$ | 23.6 | 7.77 | 18.4 | 288 | 4.86 $\times 10^{-3}$ | 437 |
| $\Phi_{(R,n_6)}$ | 0.484 | 1.00 $\times 10^{-3}$ | 9.30 | 1.00 $\times 10^{-3}$ | 21.8 | 8.57 | 22.4 | 312 | 4.85 $\times 10^{-3}$ | 439 |
| $\Phi_{(R,n_7)}$ | 0.524 | 5.43 $\times 10^{-3}$ | 9.16 | 3.10 $\times 10^{-3}$ | 24.0 | 9.40 | 22.3 | 311 | 4.86 $\times 10^{-3}$ | 448 |
| $\Phi_{(R,n_8)}$ | 0.518 | 4.29 $\times 10^{-3}$ | 9.56 | 1.41 $\times 10^{-2}$ | 24.0 | 9.40 | 21.0 | 316 | 4.62 $\times 10^{-3}$ | 455 |
| $\Phi_{optimal}$ (= $\Phi_{(R,44)}$) | 0.529 | 5.99 $\times 10^{-3}$ | 8.95 | 1.95 $\times 10^{-2}$ | 20.9 | 9.40 | 18.8 | 305 | 4.82 $\times 10^{-3}$ | 453 |

**Supplementary Table S4. Evolution of the model parameters during the route optimization (3 significant digits)**

*The model parameter set to be optimized is $\Phi = \{w_{ee}, w_{ie}, w_{ei}, w_{ii}, \tau_i, \tau_e, r_{E,b}, c, s, k\}$ (Equation (1)). During the route optimization, the network model evolution process is from $\Phi_{ini}$ (the initial parameter set) to $\Phi_{optimal}$ (the optimal parameter set). $\Phi_{pre}$ is the result of the preliminary fit (**Supplementary Notes**). $\Phi_{(G,\bullet)}$ (respectively, $\Phi_{(R,\bullet)}$) represents that it is a parameter set in Global Stage (respectively, Refining Stage) of the route optimization. $\Phi_{Hebbian}$ (= $\Phi_{(G,1)}$) is the output of 1st optimization iteration starting from $\Phi_{pre}$ (**Fig. 3**, **Fig. 4C**), and it represents a typical Hebbian network with strong recurrent excitation and weak inhibition (**Results**). The subscripts $n_2$ to $n_8$ represent the selected sample indices during the route optimization, $n_2 = 2$, $n_3 = 3$, $n_4 = 4$, $n_5 = 5$, $n_6 = 6$, $n_7 = 21$, and $n_8 = 42$. For example, $n_7 = 21$ means that $\Phi_{(R,n_7)}$ is the output of the 21 st optimization iteration during the optimization main stages (Global Stage and Refining Stage) starting from $\Phi_{pre}$ . $\Phi_{pre} \rightarrow \Phi_{(G,n_4)}$ is Global Stage of the route optimization, and $\Phi_{(G,n_4)} \rightarrow \Phi_{optimal}$ is Refining Stage of the route optimization (**Supplementary Notes**). $\Phi_{optimal}$ is the output of the 44th optimization iteration starting from $\Phi_{pre}$ . The final output $\Phi_{optimal}$ (**Fig. 3**, **Fig. 4C**) represents a typical Balanced Amplification (BA) network with balanced recurrent excitation and feedback inhibition (**Results**). The networks related to the 9 parameter sets from $\Phi_{Hebbian}$ to $\Phi_{optimal}$ are illustrated in **Fig. 4**.*

| | ER (total fitting error) | NMSE (5~50Hz) | NMSE (100Hz) | NMSE (200Hz) |
|---|---|---|---|---|
| $\Phi_{ini}$ | >100% | >100% | >100% | >100% |
| $\Phi_{pre}$ | 41.59% | 17.02% | 27.79% | 79.95% |
| $\Phi_{Hebbian}$ (= $\Phi_{(G,1)}$) | 35.92% | 17.75% | 23.27% | 66.75% |
| $\Phi_{(G,n_2)}$ | 26.60% | 14.00% | 18.17% | 47.63% |
| $\Phi_{(G,n_3)}$ | 17.17% | 13.68% | 8.20% | 29.64% |
| $\Phi_{(G,n_4)}$ | 9.54% | 13.72% | 4.29% | 10.62% |
| $\Phi_{(R,n_5)}$ | 7.90% | 14.43% | 3.58% | 5.70% |
| $\Phi_{(R,n_6)}$ | 7.84% | 14.21% | 3.62% | 5.68% |
| $\Phi_{(R,n_7)}$ | 7.78% | 13.74% | 3.37% | 6.23% |
| $\Phi_{(R,n_8)}$ | 7.70% | 13.97% | 3.45% | 5.68% |
| $\Phi_{optimal}$ (= $\Phi_{(R,44)}$) | 7.62% | 13.89% | 3.24% | 5.73% |

#### Supplementary Table S5. Evolution of the model fitting errors during the route optimization

*See the caption of **Supplementary Table S4** for the meanings of the model parameter sets from $\Phi_{ini}$ to $\Phi_{optimal}$. "ER" represents the total fitting error computed based on data across varying DBS frequencies (5~200 Hz) (Equation (2)). "NMSE" represents the "normalized mean squared error". "NMSE(5~50 Hz)" represents the model fit NMSE computed based on the concatenated data from DBS frequencies 5~50 Hz. NMSE(100 Hz) and NMSE(200 Hz) are computed with data from the 100Hz and 200Hz DBS, respectively. Note that the route optimization reduces the fitting errors fast, and the errors are already small for the model parameter set $\Phi_{(R,n_5)}$, which is the output of the 5$^{th}$ optimization iteration ($n_5 = 5$) starting from $\Phi_{pre}$ (caption of **Supplementary Table S4**). In order to further decrease the fitting errors so that the optimization exit rule (Equation (22) in **Supplementary Notes**) is satisfied, optimization iterations continued until we obtain $\Phi_{optimal}$, which is the output of the 44$^{th}$ optimization iteration starting from $\Phi_{pre}$. The networks related to the 9 parameter sets from $\Phi_{Hebbian}$ to $\Phi_{optimal}$ are illustrated in **Fig. 4**.*

| | $S = \begin{pmatrix} s_{D\xi} & s_{DI} \\ s_{I\xi} & s_{II} \end{pmatrix}$ | $\rho_{inh,D}$ | $\lambda_1$ | $\boldsymbol{v_1} = \begin{pmatrix} v_{1,1} \\ v_{2,1} \end{pmatrix}$ | $\lambda_2$ | $\boldsymbol{v_2} = \begin{pmatrix} v_{1,2} \\ v_{2,2} \end{pmatrix}$ |
|---|---|---|---|---|---|---|
| $\Phi_{Hebbian}$ $(= \Phi_{(G,1)})$ | $\begin{pmatrix} 34.9 & 3.58 \\ 0.372 & 0.688 \end{pmatrix}$ | 0.103 | 34.9 | $\begin{pmatrix} 1.00 \\ 1.09 \times 10^{-2} \end{pmatrix}$ | 0.649 | $\begin{pmatrix} -0.104 \\ 0.995 \end{pmatrix}$ |
| $\Phi_{(G,n_2)}$ | $\begin{pmatrix} 52.3 & 12.4 \\ 0.135 & 5.25 \times 10^{-3} \end{pmatrix}$ | 0.237 | 52.3 | $\begin{pmatrix} 1.00 \\ 2.57 \times 10^{-3} \end{pmatrix}$ | $-2.66 \times 10^{-2}$ | $\begin{pmatrix} -0.230 \\ 0.973 \end{pmatrix}$ |
| $\Phi_{(G,n_3)}$ | $\begin{pmatrix} 59.2 & 18.5 \\ 0.174 & 5.54 \times 10^{-3} \end{pmatrix}$ | 0.313 | 59.2 | $\begin{pmatrix} 1.00 \\ 2.93 \times 10^{-3} \end{pmatrix}$ | $-4.88 \times 10^{-2}$ | $\begin{pmatrix} -0.299 \\ 0.954 \end{pmatrix}$ |
| $\Phi_{(G,n_4)}$ | $\begin{pmatrix} 52.9 & 32.7 \\ 0.186 & 5.19 \times 10^{-3} \end{pmatrix}$ | 0.618 | 53.0 | $\begin{pmatrix} 1.00 \\ 3.52 \times 10^{-3} \end{pmatrix}$ | $-0.110$ | $\begin{pmatrix} -0.525 \\ 0.851 \end{pmatrix}$ |
| $\Phi_{(R,n_5)}$ | $\begin{pmatrix} 50.0 & 44.2 \\ 0.103 & 5.10 \times 10^{-3} \end{pmatrix}$ | 0.883 | 50.1 | $\begin{pmatrix} 1.00 \\ 2.05 \times 10^{-3} \end{pmatrix}$ | $-8.55 \times 10^{-2}$ | $\begin{pmatrix} -0.661 \\ 0.750 \end{pmatrix}$ |
| $\Phi_{(R,n_6)}$ | $\begin{pmatrix} 51.1 & 47.4 \\ 0.106 & 5.10 \times 10^{-3} \end{pmatrix}$ | 0.928 | 51.2 | $\begin{pmatrix} 1.00 \\ 2.06 \times 10^{-3} \end{pmatrix}$ | $-9.26 \times 10^{-2}$ | $\begin{pmatrix} -0.680 \\ 0.734 \end{pmatrix}$ |
| $\Phi_{(R,n_7)}$ | $\begin{pmatrix} 55.4 & 50.8 \\ 0.574 & 1.72 \times 10^{-2} \end{pmatrix}$ | 0.918 | 55.9 | $\begin{pmatrix} 1.00 \\ 1.03 \times 10^{-2} \end{pmatrix}$ | $-0.505$ | $\begin{pmatrix} -0.673 \\ 0.740 \end{pmatrix}$ |
| $\Phi_{(R,n_8)}$ | $\begin{pmatrix} 52.7 & 51.2 \\ 0.437 & 7.56 \times 10^{-2} \end{pmatrix}$ | 0.972 | 53.1 | $\begin{pmatrix} 1.00 \\ 8.23 \times 10^{-3} \end{pmatrix}$ | $-0.346$ | $\begin{pmatrix} -0.695 \\ 0.719 \end{pmatrix}$ |
| $\Phi_{optimal}$ $(= \Phi_{(R,44)})$ | $\begin{pmatrix} 54.4 & 49.3 \\ 0.616 & 0.107 \end{pmatrix}$ | 0.906 | 54.9 | $\begin{pmatrix} 1.00 \\ 1.12 \times 10^{-2} \end{pmatrix}$ | $-0.446$ | $\begin{pmatrix} -0.668 \\ 0.744 \end{pmatrix}$ |

**Supplementary Table S6. Evolution of the effective input matrix during the physiological range of route optimization (3 significant digits)**

*We show the effective input matrices in the physiological range (**Fig. 3A**), with total fitting error (ER) < 40% (**Supplementary Table S5**); we focus on the network models in this range to effectively interpret the underlying physiological meanings. See the caption of **Supplementary Table S4** for the meanings of the model parameter sets from $\Phi_{Hebbian}$ to $\Phi_{optimal}$. "S" is the effective input matrix (Equation (3)); $(\lambda_i, \boldsymbol{v_i})$ is the $i^{th}$ eigenpair of S, i.e., the $i^{th}$ eigenvalue and the associated eigenvector. "$\rho_{inh,D}$" is the inhibition strength ratio, which is the ratio of inhibitory to excitatory effective inputs into the neurons in the ventral intermediate nucleus (Vim) directly receiving DBS (Equation (4)). The networks related to the 9 parameter sets from $\Phi_{Hebbian}$ to $\Phi_{optimal}$ are illustrated in **Fig. 4**.*

| DBS frequency / rate model | high frequency | | low frequency | | | | |
|---|---|---|---|---|---|---|---|
| | 100Hz | 200Hz | 5Hz | 10Hz | 20Hz | 30Hz | 50Hz |
| **single-ensemble (Vim)** | 25.00% | 54.40% | 15.13% | 38.12% | 17.11% | 12.64% | 15.91% |
| **Hebbian network** | 23.27% | 66.75% | 18.11% | 39.75% | 15.04% | 9.03% | 11.07% |
| **Balanced Amplification network** | 3.24% | 5.73% | 14.46% | 27.53% | 16.13% | 8.26% | 4.59% |

#### Supplementary Table S7. NMSE of the rate model fits to experimental data from each DBS frequency

*We compute the normalized mean squared error (NMSE) of the rate model fits to Vim-DBS experimental data. We show the NMSE of the model fit to data from each DBS frequency (5~200 Hz), and compare 3 rate models: the single-ensemble model of the Vim neural group directly receiving DBS [2], the rate network model with the Hebbian mechanism, and the rate network model with the Balanced Amplification (BA) mechanism.*

| | N | a | b | c | d | $I_{bias}$ | SD($\xi$) | k | $\beta$ | $\theta$ | SD($\zeta$) |
|---|---|---|---|---|---|---|---|---|---|---|---|
| **Vim** | 20 | 0.02 | 0.25 | -65 | 0.05 | 0.45 | 0.225 | 1/40 | 12 | 30 | 0.225 |
| **Cerebellum** | 100 | 0.02 | 0.2 | -65 | 8 | 6.5 | 3.25 | $w_{ee}/700$ | | | 3.25 |
| **TRN** | 40 | 0.015 | 0.25 | -65 | 2.05 | 0.45 | 0.225 | $w_{ie}/700$ | | | 0.225 |

#### Supplementary Table S8. Parameters in the Izhikevich spiking network model

*We show the Izhikevich spiking network model parameters. "Vim" is the ventral intermediate nucleus, and "TRN" is the thalamic reticular nucleus. "N" represents the number of neurons in each group. {a, b, c, d} are the Izhikevich model parameters consistent with previous works. $I_{bias}$ is the biased current injected to each neuron in a group, and SD($\xi$) is the standard deviation of the Gaussian white noise current ($\xi$) injected to the neuron. "k" is the scaling parameter of the DBS-induced post-synaptic current modeled by the Tsodyks & Markram model [18,21]. "$w_{ee}$" represents the connectivity strength from Vim efferent to cerebellum, and "$w_{ie}$" represents the connectivity strength from Vim efferent to TRN (Equation (22)). $w_{ee}$ and $w_{ie}$ are from **Supplementary Table S4**, in which we focus on the two network mechanisms – Hebbian ($\Phi_{Hebbian}$) and Balanced Amplification ($\Phi_{optimal}$). "$\beta$" is a scaling parameter of the effect of synaptic connections within the network. "$\theta$" is the neuronal mean firing threshold, and SD($\zeta$) is the standard deviation of the threshold fluctuation $\zeta$, which is modeled by Gaussian white noise.*

| u | U | x | $\tau_f$ | $\tau_r$ |
|---|---|---|---|---|
| 0.35 | 0.5 | 1 | 100 | 100 |

#### Supplementary Table S9. Tsodyks & Markram model parameters used in the Izhikevich spiking network model

*We show the parameters of the Tsodyks & Markram (TM) model [21,22] of the synapses used in the Izhikevich spiking network model. This TM model is simulated by the built-in function "tsodyks2_synapse" of Neural Simulation Tool (NEST) platform [23], and the simulation result (**Supplementary Fig. S9**) is consistent with the TM model (**Supplementary Table S2**) used in our rate model (**Supplementary Fig. S2**). "u" is the probability of* *neurotransmitter release. "U" is the parameter determining the increase in u with each spike. "x" is a scaling parameter of the amount of neurotransmitter release. "$\tau_f$" is the time constant (ms) for facilitation of release, and "$\tau_r$" is the time constant (ms) for recovery (returning the neurotransmitters to the available pool)* [21].

| length of initial transient period (ms) | category (DBS frequency) |
|---|---|
| 705.2 | 100 Hz |
| 917.2 | 100 Hz |
| 743.7 | 100 Hz |
| 284.9 | 100 Hz |
| 448.2 | 100 Hz |
| 883.4 | 100 Hz |
| 808.6 | 100 Hz |
| 734.9 | 100 Hz |
| 228.7 | 200 Hz |
| 207.2 | 200 Hz |
| 349.2 | 200 Hz |
| 210.9 | 200 Hz |
| 278.4 | 200 Hz |

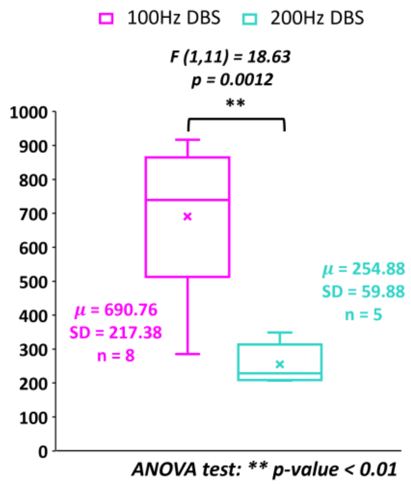

#### Supplementary Table S10. Length of initial transient responses in clinical recordings during high-frequency DBS

*The clinical data include 8 recordings during 100-Hz DBS and 5 recordings during 200-Hz DBS. For each recording, the length of the initial transient response is computed as the distance from time 0 to the last spike timing during the initial transient period with a large firing rate. For 100-Hz and 200-Hz DBS, the statistics are 690.76 $\pm$ 217.38 ms and 254.88 $\pm$ 59.88 ms (mean $\pm$ standard deviation), respectively. The transient response length during 100-Hz DBS is significantly longer than that during 200-Hz DBS (ANOVA, F (1,11) = 18.63, p = 0.0012).*

**100Hz DBS (pulse #50-52):**

| neuron / Pulse id | Vim 03 (row 1) | | | Vim 19 (row 2) | | | Vim 10 (row 3) | | |
|---|---|---|---|---|---|---|---|---|---|
| | peak | valley | **diff** | peak | valley | **diff** | peak | valley | **diff** |
| #50 | 0.259 | 0.234 | **0.025** | 0.210 | -0.132 | **0.342** | 0.510 | 0.364 | **0.146** |
| #51 | 0.151 | 0.139 | **0.012** | 0.442 | 0.271 | **0.171** | 0.388 | 0.239 | **0.149** |
| #52 | 0.315 | 0.273 | **0.042** | 1.917 | 1.680 | **0.237** | 0.425 | 0.237 | **0.188** |

**100Hz DBS (pulse #250-252):**

| neuron / Pulse id | Vim 03 (row 1) | | | Vim 19 (row 2) | | | Vim 10 (row 3) | | |
|---|---|---|---|---|---|---|---|---|---|
| | peak | valley | **diff** | peak | valley | **diff** | peak | valley | **diff** |
| #250 | 0.205 | 0.176 | **0.029** | 1.575 | 0.308 | **1.267** | 0.308 | 0.212 | **0.096** |
| #251 | 0.232 | 0.205 | **0.027** | 1.658 | 0.415 | **1.243** | 0.442 | 0.295 | **0.147** |
| #252 | 0.193 | 0.134 | **0.059** | 1.387 | 0.129 | **1.258** | 0.364 | 0.100 | **0.264** |

**100Hz DBS (pulse #279-281):**

| neuron / Pulse id | Vim 03 (row 1) | | | Vim 19 (row 2) | | | Vim 10 (row 3) | | |
|---|---|---|---|---|---|---|---|---|---|
| | peak | valley | **diff** | peak | valley | **diff** | peak | valley | **diff** |
| #279 | 0.156 | 0.078 | **0.078** | 1.260 | 0.542 | **0.718** | 0.637 | 0.488 | **0.149** |
| #280 | 0.134 | 0.085 | **0.049** | 1.567 | 0.715 | **0.852** | 0.557 | 0.469 | **0.088** |
| #281 | 0.203 | 0.171 | **0.032** | 1.677 | 0.779 | **0.898** | 0.640 | 0.627 | **0.013** |

**200Hz DBS (pulse #50-52):**

| neuron / Pulse id | Vim 10 (row 1) | | | Vim 11 (row 2) | | | Vim 14 (row 3) | | |
|---|---|---|---|---|---|---|---|---|---|
| | peak | valley | **diff** | peak | valley | **diff** | peak | valley | **diff** |
| #50 | 0.486 | 0.408 | **0.078** | 0.159 | 0.154 | **0.005** | 0.562 | 0.469 | **0.093** |
| #51 | 0.481 | 0.396 | **0.085** | 0.254 | 0.200 | **0.054** | 0.579 | 0.496 | **0.083** |
| #52 | 0.481 | 0.383 | **0.098** | 0.305 | 0.244 | **0.061** | 0.527 | 0.503 | **0.024** |

**200Hz DBS (pulse #250-252):**

| neuron / Pulse id | Vim 10 (row 1) | | | Vim 11 (row 2) | | | Vim 14 (row 3) | | |
|---|---|---|---|---|---|---|---|---|---|
| | peak | valley | **diff** | peak | valley | **diff** | peak | valley | **diff** |
| #250 | 0.376 | 0.168 | **0.208** | 0.239 | 0.164 | **0.075** | 0.281 | 0.251 | **0.03** |
| #251 | 0.437 | 0.195 | **0.242** | 0.271 | 0.193 | **0.078** | 0.295 | 0.239 | **0.056** |
| #252 | 0.386 | 0.205 | **0.181** | 0.254 | 0.164 | **0.09** | 0.330 | 0.283 | **0.047** |

**200Hz DBS (pulse #450-452):**

| neuron / Pulse id | Vim 10 (row 1) | | | Vim 11 (row 2) | | | Vim 14 (row 3) | | |
|---|---|---|---|---|---|---|---|---|---|
| | peak | valley | **diff** | peak | valley | **diff** | peak | valley | **diff** |
| #450 | 0.271 | 0.215 | **0.056** | 0.254 | 0.114 | **0.140** | 0.491 | 0.444 | **0.047** |
| #451 | 0.332 | 0.225 | **0.107** | 0.273 | 0.120 | **0.153** | 0.508 | 0.481 | **0.027** |
| #452 | 0.295 | 0.232 | **0.063** | 0.305 | 0.198 | **0.107** | - | - | - |

#### Supplementary Table S11. Biomarker of inhibition in single-unit membrane potential data recorded during high frequencies of DBS

*As shown in **Fig. 8**, the biomarker of inhibition is characterized by the difference between the "peak" of hyperpolarization and a preceding "valley" representing the initialization of inhibition. "row 1" indicates the data corresponding to the 1st row in **Fig. 8**; similar meanings for "row 2" and "row 3". The data "Vim 200Hz, neuron Vim 14, pulse #452" is excluded in the computation and statistical test, because the biomarker is hidden by a spiking (depolarizing) activity (**Fig. 8**). There are 5 Vim neurons (Vim 03, 10, 11, 14, 19) (out of a total sample of 19 Vim neurons, one from each patient, see **Methods**) that exhibit the evoked inhibitory activities during high frequency DBS.*

#### Supplementary video

##### Supplementary Video S1. Evolution of network mechanisms during route optimization

*We illustrate the evolution of network mechanisms during the route optimization process. During route optimization, the network mechanisms evolve from Hebbian to Balanced Amplification. The network mechanisms are quantitatively defined by the inhibition strength ratio of the effective inputs into Vim neurons receiving DBS ($\rho_{inh,D}$, Equation (4)). In this video, we present the network model fits corresponding to 6 parameter sets (**Supplementary Table S6**): $\Phi_{Hebbian}$, $\Phi_{(G,n_2)}$, $\Phi_{(G,n_3)}$, $\Phi_{(G,n_4)}$, $\Phi_{(R,n_5)}$, and $\Phi_{optimal}$. We present the network model fits to the experimental data from 3 DBS frequencies: 5 Hz, 100 Hz and 200 Hz.*
